# Gut microbiome-dependent IL-1 signaling is a mediator of ACVR1^R206H^-driven heterotopic ossification

**DOI:** 10.64898/2026.04.05.716562

**Authors:** Hannah M. Herzog, Camille Fang, Liam Lam, Katherine Jin, Ariane Zamarioli, Ethan Dinh, Chhedi Lal Gupta, Aditi Sharma, Tania Moody, Jessica L. Pierce, Michael S. Hohl, Sarah W. Takimoto, Svetlana Lyalina, Kelly L. Wentworth, Kristie Yu, Vivian F. Lu, Isadora Mamikunian, Natasha K. Hunt, Susan Lynch, Katherine S. Pollard, Christopher J. Hernandez, Daniel S. Perrien, Edward C. Hsiao

## Abstract

Inflammatory diseases cause significant morbidity and mortality, but their pathobiology is often difficult to dissect due to complex genetic-environmental interactions. Genetic forms of heterotopic ossification, such as fibrodysplasia ossificans progressiva (FOP), reduce genetic variability, allowing careful dissection of non-genetic drivers of inflammation. While >95% of FOP patients harbor the *ACVR1^R206H^* mutation, patients exhibit significant variability in disease progression, suggesting a role of environmental drivers. Here, we identify the gut microbiome as a regulator of inflammation-driven HO in FOP. Metagenomic profiling of cohabitating FOP/unaffected sibling pairs revealed a pathogenic gut microbiome profile in FOP patients (Bray-Curtis, p < 0.05). In *Pdgfrα-Cre/Acvr1^R206H^* (FOP) mice, gut microbiome ablation by antibiotics reduced spontaneous HO formation (47.4% reduction, p < 0.05) and reduced plasma IL-1 pathway activity. IL-1β blockade in FOP mice suppressed trauma-induced HO formation. These findings identify a gut microbiome-IL-1-HO axis with modifiable targets for developing treatments for HO and related inflammatory conditions.

**One Sentence Summary:** Antibiotic disruption of the gut microbiome reduces HO in FOP mice via an IL-1 mediated pathway.

## INTRODUCTION

Chronic inflammatory diseases are a major contributor to U.S. healthcare spending, accounting for nearly 90% of annual expenditures^1–3^. Approximately 35% of U.S. adults exhibit evidence of systemic inflammation, and relapsing/remitting inflammatory diseases such as systemic lupus erythematosus, inflammatory bowel disease (IBD), multiple sclerosis, and rheumatoid arthritis contribute disproportionately to healthcare use and economic burden^4,5^. Together, chronic inflammatory conditions account for at least $90 billion annually in direct healthcare costs and $41.2 billion in productivity losses, highlighting the need to understand disease pathobiology and identify effective management strategies^4,6^.

Many inflammatory diseases are characterized by recurrent episodes of acute symptom worsening, referred to as flares, which are periods of heightened immune activation and tissue inflammation, separated by intervals of relative quiescence. These flare-driven conditions are often multigenic and unpredictable, making it difficult to isolate environmental contributors in genetically heterogeneous human populations. Fibrodysplasia ossificans progressiva (FOP) is a rare, monogenic musculoskeletal disorder characterized by the episodic pathogenic formation of bone in soft tissues, termed heterotopic ossification (HO)^7,8^. Progressive HO leads to ankylosis of major joints, severe restriction of movement, and ultimately early mortality^9^. HO formation in FOP is often preceded by cyclic inflammatory flare-ups, but can also occur in the absence of an obvious trigger. These flares are often initiated by minor trauma in local tissue (acute or injury-induced HO)^8^ and can persist for days to months. Anti-inflammatory treatments such as glucocorticoids are the mainstay of care to limit inflammation and HO formation^10^, but are largely ineffective and unsuitable for long-term use.

Despite >95% of FOP patients carrying the recurrent c.617G>A (p.R206H) activating mutation in the Activin A Receptor, Type I (*ACVR1/ALK2*) gene^11,12^, clinical severity varies widely, with some patients becoming immobile in adolescence while others remain mobile into adulthood^8,12^. The clinical variability of FOP patients suggests that non-genetic factors, including environmental influences, modulate HO progression. In both genetic and acquired HO, ossification is driven by inflammatory responses involving the recruitment of macrophages and the release of pro-inflammatory and pro-fibrotic cytokines^13–18^, with NF-κB signaling playing a central role in regulating macrophage activity in FOP^18,19^. Non-genetic HO (e.g., post-arthroplasty) also exhibits substantial variability, lacks clear genetic drivers, and shares similar inflammatory features to FOP^20^.

Given the key role of inflammation in HO ^19–22^, the gut microbiome is of particular interest as a regulator of inflammatory signaling and mediator of environmental influences^23^. The microbiome has been linked to systemic inflammatory diseases, including rheumatoid arthritis, cardiovascular disease, and inflammatory bowel disease^24–29^, and can modulate NF-κB signaling, macrophage activity, and pro-inflammatory cytokine production^30,31^. Furthermore, microbiome-derived signals influence bone formation and function^32–34^, suggesting that the gut microbiome may contribute to osteogenesis and HO pathogenesis. Therefore, we hypothesized that the gut microbiome promotes inflammation-mediated osteogenesis in FOP and that gut microbiome ablation (GMA) would reduce new HO formation. Our findings identify the gut microbiome and IL-1 pathways as modifiable drivers of HO and suggest that targeting microbiome-immune interactions may represent a potential therapeutic strategy for FOP and other inflammation-driven disorders of ectopic ossification.

## RESULTS

### Demographic characteristics of subjects with FOP and their family controls

Patients with FOP have reported a significant frequency of gastrointestinal symptoms^35^. To better understand these symptoms, we recruited 16 subjects with FOP and 15 non-FOP siblings (**Figure 1A**) using protocols and surveys approved by the University of California at San Francisco Institutional Review Board (UCSF IRB #16-19534). **Table 1** summarizes the baseline demographic characteristics of the studied participants. The majority of participants were male (54.8%) and non-Hispanic white (64.5%) with an average age of 16.3 (± 11.3 years, mean ±SD). The average CAJIS score of patients with FOP was 9.5 (± 7, mean ±SD, maximum of 30), consistent with prior reported CAJIS scores for this age ^36^.

**Figure 1.**
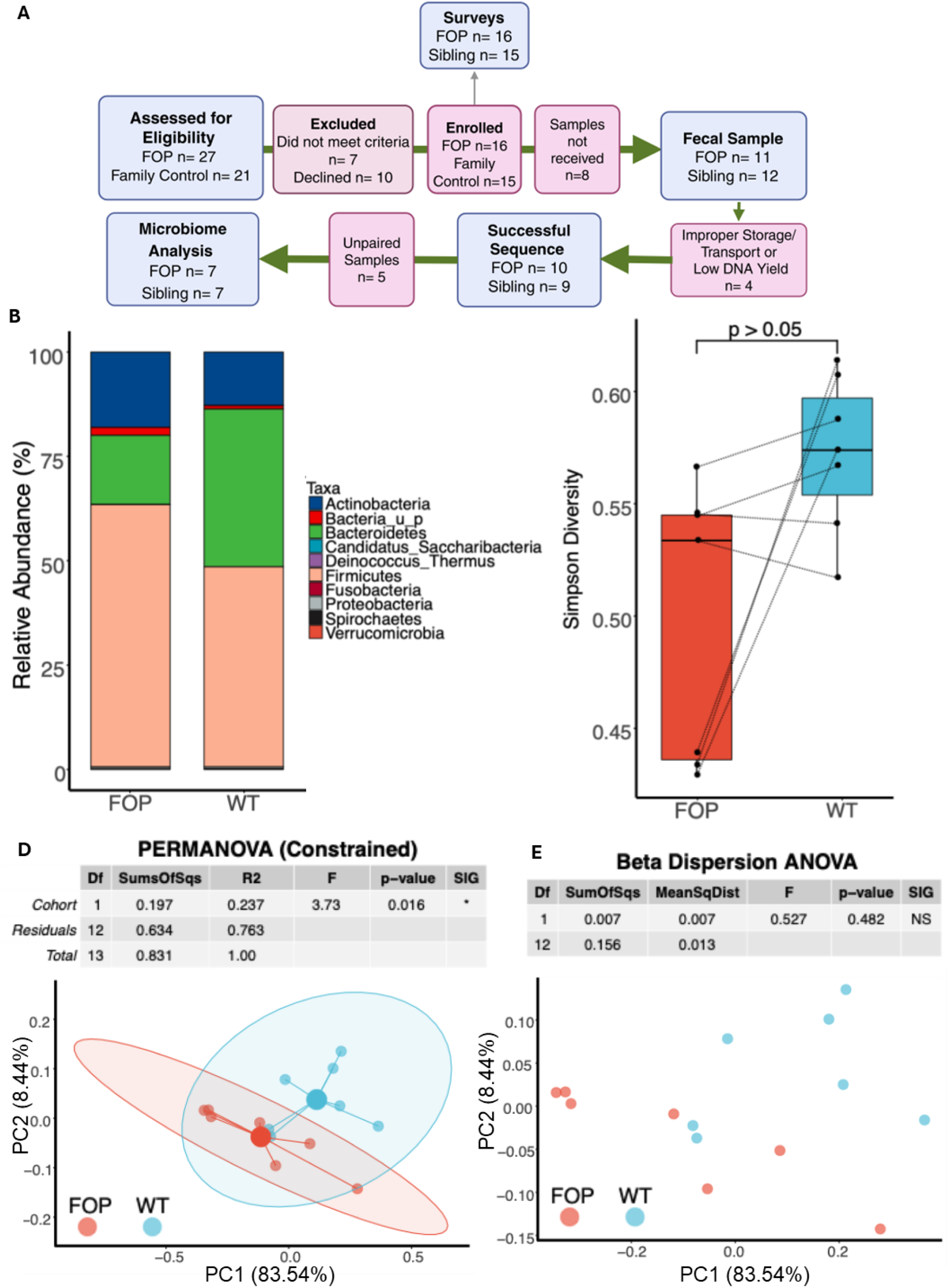
Human microbiome study design and compositional analyses (sibling-paired cohort) **(A) Study design and cohort selection.** Flow diagram of participant enrollment and sample inclusion. Sixteen individuals with fibrodysplasia ossificans progressiva (FOP) and 15 family controls were analyzed for the survey data. Stool samples were self-collected, frozen, and processed for shotgun metagenomic sequencing. Samples were excluded due to incomplete collection, low DNA yield, or lack of matched pairs, resulting in 7 sibling-paired FOP–control samples included in microbiome analyses. Created with Biorender.com. **(B) Taxonomic composition.** Stacked bar plots showing relative abundance of major microbial taxa in FOP and matched sibling control samples. Taxonomic profiling was performed using the CosmosID platform from shotgun metagenomic data. **(C) Alpha diversity.** Simpson diversity index comparing FOP and matched sibling control samples. Diversity metrics were calculated in R using the vegan package, and paired differences were assessed using Wilcoxon rank-sum tests (p > 0.05). **(D) Beta diversity (PCoA with PERMANOVA).** Principal coordinates analysis (PCoA) based on Bray–Curtis dissimilarity showing differences in overall microbiome composition between FOP and matched sibling controls. Group differences were assessed using PERMANOVA (adonis2). **(E) Beta dispersion.** Analysis of multivariate dispersion demonstrates no significant difference in within-group variability between FOP and matched sibling control samples (ANOVA, p = 0.482).

**Table 1.** Demographic and clinical characteristics of the human study cohort. The study included 31 participants, comprising 16 individuals with FOP and 15 healthy family controls. All FOP participants met clinical diagnostic criteria for classical FOP. Family controls were biological siblings living in the same household, except for one spouse control subject. Demographic information, including age, sex, and race/ethnicity, is shown for the total cohort and by group. Cumulative joint involvement scale (CAJIS) scores are presented for FOP participants from the full survey cohort and the subset included in microbiome analyses. One participant was unable to complete CAJIS assessment.

|  | Total |  | Family |  | FOP |  |
| --- | --- | --- | --- | --- | --- | --- |
| **<u>Characteristic</u>** | N | % | N | % | N | % |
| All Patients | 31 | 100.00 | 15 | 48.39 | 16 | 51.61 |
| **Age (mean, sd)** | 16.3, (11.3) | - | 16.1, (11.9) | - | 16.5, (11.1) | - |
| **Sex** |  |  |  |  |  |  |
| Male | 17 | 54.80 | 7 | 46.70 | 10 | 62.50 |
| Female | 14 | 45.20 | 8 | 53.30 | 6 | 37.50 |
| **Race/Ethnicity** |  |  |  |  |  |  |
| Non-Hispanic White | 20 | 64.50 | 9 | 60.00 | 11 | 68.80 |
| Asian | 3 | 9.70 | 2 | 13.30 | 1 | 6.20 |
| Hispanic/LatinX | 3 | 9.70 | 1 | 6.70 | 2 | 12.50 |
| Other | 2 | 6.50 | 1 | 6.70 | 1 | 6.20 |
| Decline to State | 3 | 9.70 | 2 | 13.30 | 1 | 6.20 |
|  | **FOP Total** |  | **CAJIS** |  | **Jaw CAJIS** |  |
| **<u>Study</u>** | N | % | Mean | SD | Mean | SD |
| Survey | 16 | 100 | 9.5 | 7.0 | 0.46 | 0.74 |
| Microbiome | 7 | 43.75 | 8 | 5.97 | 0.29 | 0.49 |

### Eating and clinical GI symptoms are not significantly different between patients with FOP and family controls

Clinical surveys on our cohort showed no significant differences in diet, food preferences, or appetite between patients with FOP and their siblings (**Supplemental Table 1**). Patients with FOP were more likely to avoid foods that required chewing (Fisher’s exact test, p = 0.02), consistent with the known jaw restrictions that can occur in FOP (**Table 2**). Jaw involvement was observed in 5 patients, and the mean jaw-specific CAJIS across all patients was 0.46 (scale 0-2) (**Table 1**). Patients with FOP reported more GI symptoms on average than sibling controls (**Supplemental Table 2**); however, the differences did not reach significance except for vomiting and “vomiting until stomach was empty” (Fisher’s exact test, p = 0.01 and p = 0.04, respectively) (**Table 2**). Patients with FOP more frequently reported passing urine or stool instead of gas compared to unaffected sibling controls (Fisher’s exact test, p = 0.01) (**Table 2**). This may reflect altered pelvic floor control or urgency related to musculoskeletal restriction, rather than primary gastrointestinal pathology.

**Table 2.** Significant differences in gastrointestinal symptoms and food preferences between FOP patients and family controls. Survey data were collected using REDCap-based questionnaires adapted from validated instruments (e.g., GI-PROMISE; full analysis in Supplemental Tables 1 and 2). Responses were dichotomized, and group differences were assessed using Fisher’s exact tests in Stata (p < 0.05 considered significant). Individuals with FOP more frequently reported avoidance of foods requiring chewing, vomiting, vomiting until the stomach was empty, and passing gas with stool or urine leakage compared to family controls.

|  | Family |  | FOP |  |  |
| --- | --- | --- | --- | --- | --- |
| <u>Characteristic</u> | N | % | N | % | pvalue |
| Avoided food that requires chewing | 1 | 7 | 8 | 50 | *0.02 |
| Vomit | 2 | 13 | 10 | 63 | *0.01 |
| Vomit until stomach empty | 1 | 7 | 7 | 44 | *0.04 |
| Pass gas but stool or urine come out instead | 0 | 0 | 6 | 60 | *0.01 |

### Gut microbiome composition differs between FOP patients and their sibling controls

To control for environmental confounders, we focused on participants with FOP who had an unaffected age-similar sibling living at the same home to provide a pair-wise comparison.

Seven subject pairs were identified with high quality stool samples and used for the shotgun metagenomic sequencing analysis (**Figure 1A**). Paired analyses was used to reduce the impact of environmental variables that may confound results (**Figure 1**). FOP patients had a higher relative abundance of the phyla *Firmicutes* and *Actinobacteria*, and a lower relative abundance of the phyla *Bacteroidetes* compared to WT (**Figure 1B**). Beta diversity analysis using a Bray-Curtis dissimilarity matrix revealed a significant difference in fecal microbial community composition between FOP patients and their healthy sibling controls (PERMANOVA with permutations restricted within sibling pairs, p = 0.016) (**Figure 1D-E**). In addition, a trend towards lower microbial alpha diversity (Simpson index) was seen in the FOP group at the taxonomic level (two-sided Wilcoxon, p = 0.078) (**Figure 1C**). These findings suggest that FOP patients may harbor a significantly different gut microbiome compared to their healthy siblings.

### FOP patients show gut microbial community structure differences and an increased proportion of pathogenic bacteria

To further define the microbiome community structures, we applied the DESeq2 algorithm to test for differential abundance in bacterial groups that could be driving the separation in fecal microbiome composition observed in our paired comparisons.

FOP patients showed increased abundance of *Firmicutes* (p = 0.047) and decreased *Bacteroidetes* (p = 0.016). At the species level, FOP patients had reduced abundance of *Alistipes finegoldii*, *A. senegalensis*, *Alistipes sp. HGB5*, *Lachnospiraceae bacterium 3_1_57FAA_CT1*, *Prevotella copri*, and *Veillonella_u_s*., and increased abundance of *Clostridium innocuum*, *Bacterium LF 3*, *Erysipelotrichaceae bacterium 21_3*, *Streptococcus mitis*, *Turicibacter sanguinis*, and *Turicibacter_sp_HGF1* (**Supplemental Table 3**). Notably, taxa with immunoregulatory or anti-inflammatory associations (*A. senegalensis*, *A. finegoldii* ^37,38^, *Lachnospiraceae bacterium 3_1_57FAA_CT1* ^39,40^) were depleted, whereas inflammatory-associated taxa and pathobionts (*S. mitis* ^41,42^, *C. innocuum* ^43,44^, *Erysipelotrichaceae bacterium 21_3* ^45,46^) were enriched in FOP patients (**Supplemental Table 4**).

MetaCyc-based functional profiling combined with linear discriminant analysis effect size (LEfSe) analysis revealed significant enrichment of bacterial ppGpp biosynthesis, prokaryotic TCA cycle activity, and arginine biosynthesis pathways in FOP relative to WT microbiomes (**Supplemental Table 5**). These pathways indicate a microbiome adapted to metabolic stress and altered nutrient availability^47–49^.

Together, these compositional differences and functional features indicated that patients with FOP harbor a gut microbiota that has shifted away from anti-inflammatory or immune-supportive taxa and towards a microbiome enriched for pathways with pathogenic or pro-inflammatory characteristics.

### FOP mice with gut microbiome ablation (GMA) show decreased chronic HO volume

Because FOP is rare, human microbiome studies in genetically matched cohorts are limited. To address this, we used an established FOP mouse model^50^ with conditional expression of *ACVR1^R206H^*driven by *PDGFRα-Cre* (**Figure 2A**). These mice develop HO spontaneously and after trauma, with mutation expression restricted to fibroadipocytes, preserving a normal immune system and GI tract. This model enables direct manipulation of the gut microbiome to test its impact on HO (**Figure 2B**).

**Figure 2.**
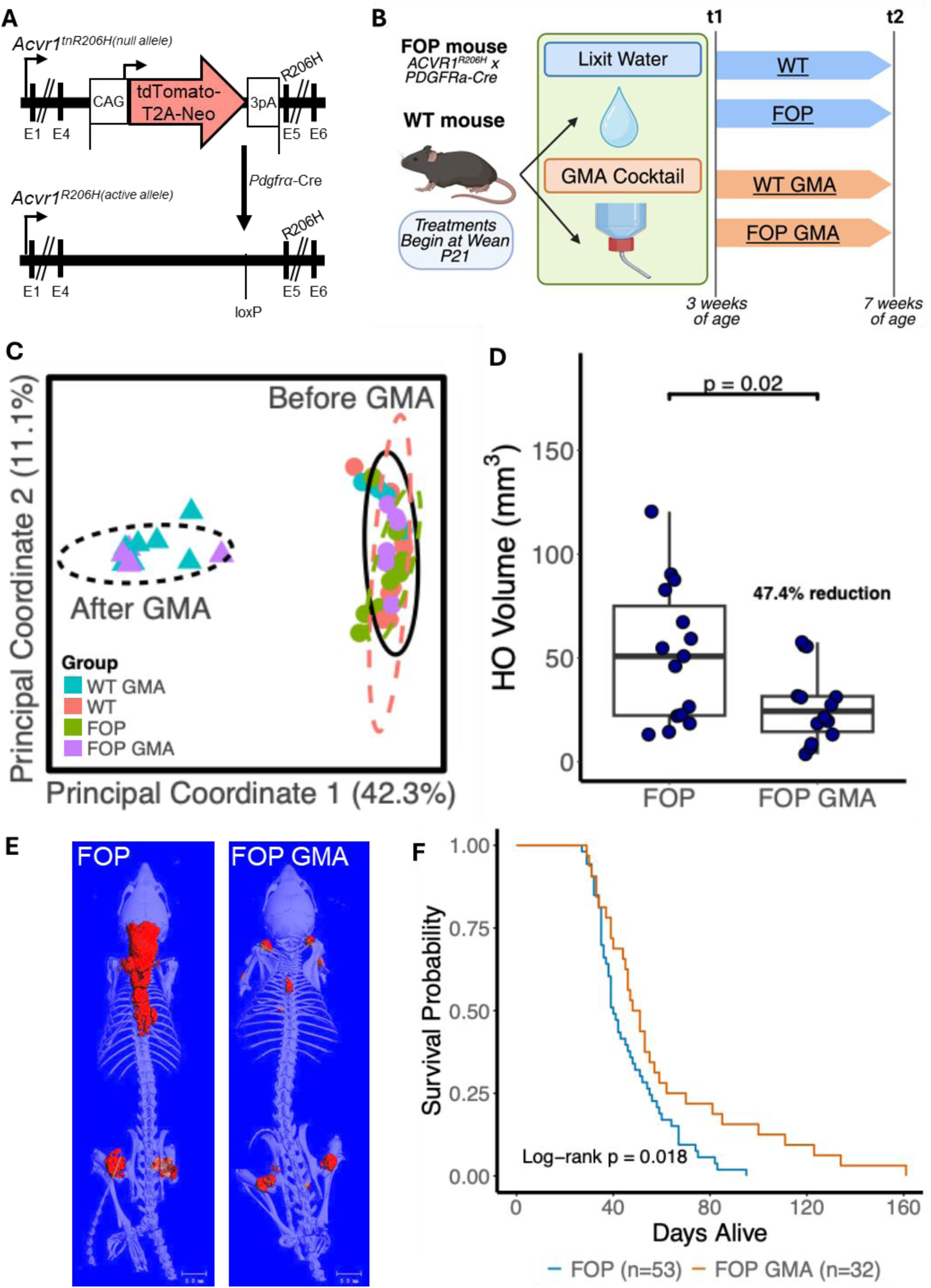
Gut Microbiome Ablation (GMA) Reduces Heterotopic Ossification and Improves Survival in FOP mice. **(A) Genetic model.** Schematic of the *PDGFRα-Cre/ACVR1^R206H^* FOP mouse model, in which expression of the constitutively active *ACVR1^R206H^*allele is expressed in fibroadipogenic progenitors following Cre recombination. **(B) Experimental design.** FOP and control (WT) littermates were treated beginning at weaning (postnatal day 21) with either normal water or a broad-spectrum antibiotic cocktail (1 g/L ampicillin + 0.5 g/L metronidazole, neomycin, vancomycin) to induce gut microbiome ablation (GMA). Samples were collected at baseline (t1) and after treatment (t2). Created with Biorender.com. **(C) Beta diversity.** Principal coordinates analysis (PCoA) of Bray–Curtis dissimilarity showing separation of microbial communities before (circles) and after GMA (triangles). Analyses were performed using phyloseq and vegan, with PERMANOVA used to assess group differences. Gut microbial composition was significantly different post-GMA (PERMANOVA, p=0.001) **(D) Heterotopic ossification (HO) volume.** MicroCT quantification of HO volume in FOP and FOP GMA mice. HO volume was significantly reduced following GMA (Welch’s t-test, p = 0.02), corresponding to a 47.4% reduction. **(E) Representative microCT imaging.** Whole-body microCT images demonstrating reduced ectopic bone formation in FOP GMA (n=15) compared to untreated FOP mice (n=12). Ectopic bone volume is shown in red, and the native skeleton in shown in white. Representative images were chosen based on the mean HO volume of each group. **(F) Survival analysis.** Kaplan–Meier survival curves comparing FOP (n=53) and FOP GMA mice (n=32). Survival was significantly improved following GMA (log-rank test, p = 0.018).

Shotgun metagenomic profiling of the mouse fecal microbiome revealed similar community diversity and structure between FOP and littermate control (WT) mice prior to intervention (timepoint t1) (**Supplemental Figure 1A**). Measures of alpha and beta diversity did not identify overt genotype- or sex-associated differences at baseline (**Figure 2C**, **Supplemental Figure 1B**), and species-level and functional analyses showed only minor variation between groups (**Supplemental Figure 2, Supplemental Table 6**). These findings indicate no major differences at baseline in the FOP vs. WT mice, consistent with *ACVR1^R206H^* expression restricted to *PDGFRα*-lineage cells and not in the gut.

To test whether depletion of the gut microbiome alters systemic inflammation and therefore HO severity in FOP, gut microbiome ablation (GMA) was induced using a broad-spectrum antibiotic cocktail administered in drinking water for four weeks beginning at postnatal day 21 (**Figure 2B**). As expected, GMA resulted in a significant change in gut microbial composition (Bray-Curtis, PERMANOVA p = 0.001) (**Figure 2C**) and a substantial depletion of taxa in both WT and FOP mice (**Supplemental Figure 2B, Supplemental Figure 3-4**).

Whole-body microCT performed at tissue collection at the endpoint (timepoint t2) revealed a marked reduction in spontaneous HO across the FOP mouse groups. Compared to untreated FOP controls (n = 15), FOP mice treated with GMA (n = 12) exhibited a 47.7% reduction in total HO volume (p = 0.02) (**Figure 2D-E**).

### FOP mice with GMA survive longer than untreated mice

The FOP mouse model used in this study develops severe spontaneous HO, which frequently leads to early mortality^51,52^. Given the efficacy of GMA in reducing spontaneous HO burden, we next asked whether GMA conferred a longitudinal survival benefit in FOP mice. To do so, we ablated the gut microbiota in FOP mice from postnatal day 21 until natural death or humane endpoint (including Body Condition Score ≤2). Survival times analyzed on a Kaplan-Meyer curve showed that FOP mice at baseline (n = 53) survived on average to postnatal day 46.5, whereas FOP mice treated with GMA (n = 32) survived significantly longer, to an average of postnatal day 60.2 (log-rank p = 0.018) (**Figure 2F**). The oldest mice showed nearly a doubling of survival (95 days vs. 161 days).

### FOP mice with GMA showed no significant differences in plasm MMP-9 levels

A prior study identified matrix metalloproteinase-9 (MMP-9) as a key inflammatory mediator linking macrophage activation to HO in FOP, with reduced MMP-9 activity attenuating disease severity in both mice and a human patient^53^. MMP-9 mutant macrophages produce less Activin-A, a key inflammatory stimulus for HO progression, and even a 50% decrease in MMP-9 activity can prevent HO formation in mice^53^. Because tetracycline antibiotics, such as minocycline, can reduce MMP-9 levels ^53–55^, we tested whether our antibiotic cocktail may also have direct effects on the plasma MMP-9 levels of FOP mice. Within our broad-spectrum antibiotic cocktail, vancomycin and neomycin are poorly absorbed through the gut and have low blood bioavailability; metronidazole and ampicillin are readily absorbed but are not known to inhibit MMP-9 activity. Plasma from FOP and control mice, +/- GMA, showed no significant differences in plasma MMP-9 levels (**Figure 3A**). This lack of variation suggests that antibiotic-mediated GMA does not affect MMP-9 activity and is unlikely to be the mechanism responsible for the reduced HO volume observed with GMA treatment.

**Figure 3:**
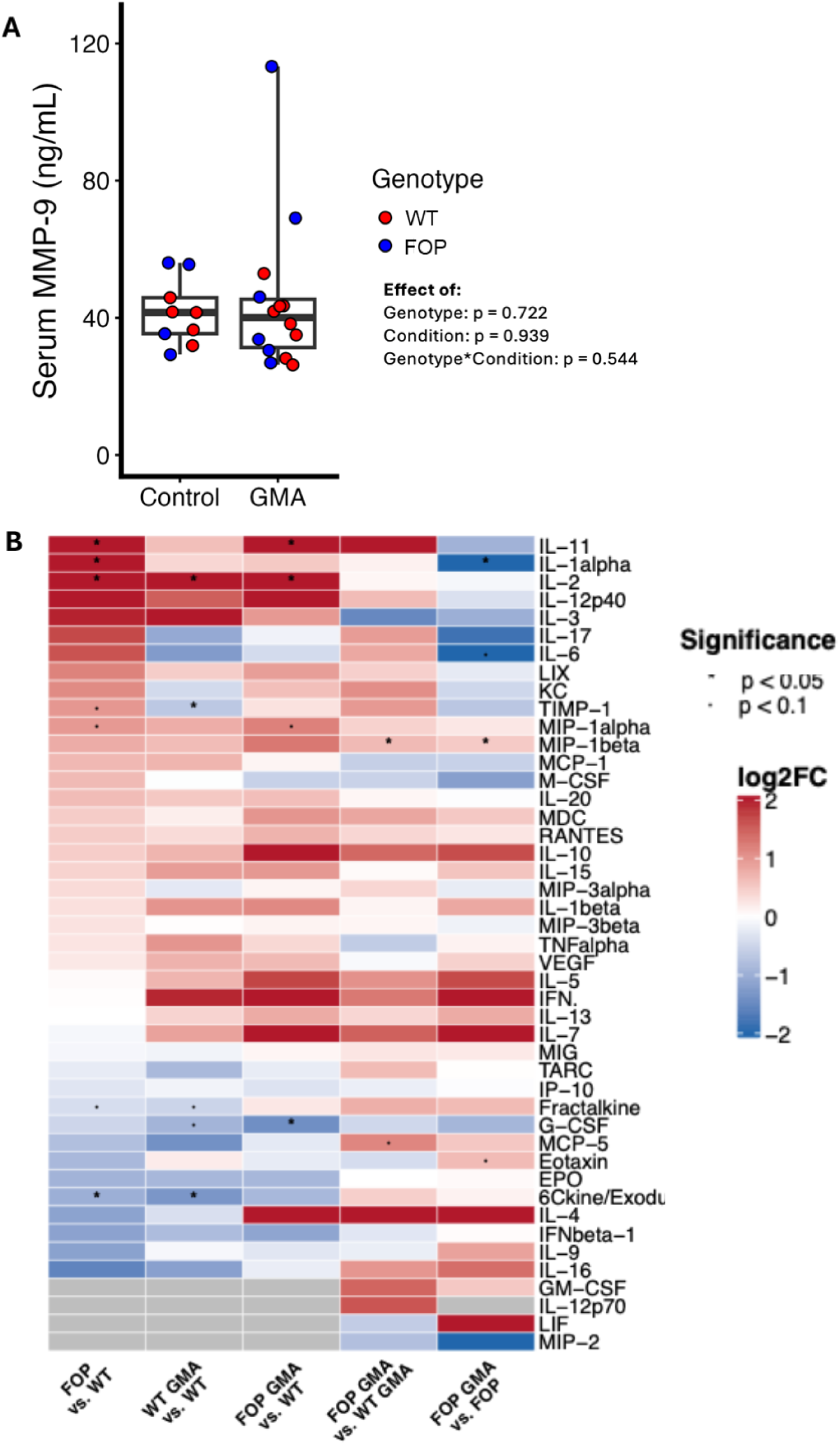
Gut Microbiome Ablation Reduces Systemic Inflammation in FOP Mice. **(A)** Serum MMP-9 concentrations in WT and FOP mice +/- GMA. Mice received a broad-spectrum antibiotic cocktail (ampicillin, neomycin, metronidazole, and vancomycin) prior to serum collection. Individual points represent biological replicates and boxes represent group distributions. Effects of genotype, treatment condition (control vs GMA), and their interaction were tested using a linear mixed-effects model. No significant effects were observed for genotype, condition, or genotype × condition interaction. **(B)** Heatmap of circulating cytokines measured in plasma using a 44-plex Luminex assay (EVE Technologies). Cytokine abundance values were aggregated by experimental group (WT, WT GMA, FOP, FOP GMA), and log₂ fold-change values were calculated relative to WT or directly comparing FOP GMA to FOP. Colors represent relative changes in cytokine abundance (log₂ fold-change scale), with red indicating increased levels and blue indicating decreased levels. Asterisks indicate cytokines with statistically significant differences between conditions (p < 0.05).

### FOP mice with GMA show suppression of inflammatory cytokines

Because MMP9 did not appear to mediate the reduction in HO observed in FOP mice following GMA treatment, we next asked whether GMA may attenuate HO by dampening systemic inflammation, which can amplify pro-osteogenic immune signaling in FOP. We quantified plasma cytokines in WT and FOP mice ± GMA. At baseline, FOP mice exhibited higher levels of multiple pro-inflammatory and pro-fibrotic mediators than WT mice, including IL-1α (p = 0.03), IL-2 (p = 0.01), MIP-1α (p = 0.06), IL-11 (p = 0.02), and TIMP-1 (p = 0.07), and lower levels of 6Ckine/Exodus 2 (p = 0.05) and fractalkine (p = 0.07) (**Figure 3B**). Several of these cytokines were previously reported to be elevated in patients with FOP or in iPSC-derived macrophages carrying the FOP mutation (**Supplemental Table 7**)^19^. Following GMA treatment, FOP mice showed decreased IL-1α (p = 0.20), TIMP-1 (p = 0.47), 6Ckine/Exodus 2 (p = 0.11), and fractalkine (p = 0.47) and were no longer different than WT mice (**Figure 3B**). In contrast, G-CSF (p = 0.03), IL-2 (p = 0.03), MIP-1α (p = 0.06), and IL-11 (p = 0.05) remained elevated post-GMA treatment in FOP compared to WT mice (**Figure 3B**). GMA treated FOP mice also exhibited reduced levels of eotaxin (p = 0.07), IL-1α (p = 0.05), IL-6 (p = 0.07), and MIP-1β (p = 0.05) compared to baseline FOP mice (**Figure 3B**).

In WT mice, GMA was associated with decreased G-CSF (p = 0.07), IL-2 (p = 0.03), and fractalkine (p = 0.07), and increased 6Ckine/Exodus 2 (p = 0.01) and TIMP-1 (p = 0.02) compared to WT at baseline (**Figure 3B**). When comparing FOP GMA to WT GMA mice, MIP-1β was significantly higher in FOP GMA mice (p = 0.05) (**Figure 3B**).

GMA reduced systemic inflammatory signaling in FOP mice by eliminating the baseline FOP versus WT differences in IL-1α, IL-2, and TIMP-1, and by reducing IL-1α (FOP GMA lower than baseline FOP) as well as eotaxin, IL-6, and MIP-1β in the FOP GMA versus baseline FOP comparison. These changes are consistent with our human datasets reporting higher IL-1α, IL-6, and/or MIP-1β in FOP serum and/or FOP iPSC-derived macrophages compared with WT controls (**Supplemental Table 7**)^16,19^.

### Antibody-mediated IL-1 blockade abolishes trauma-induced HO in FOP mice

Our GMA studies support the IL-1 pathway as a central mediator of HO in FOP. At baseline, IL-1-related components, including plasma IL-1α, were significantly elevated in FOP mice (135.1 vs. 12.6 pg/mL in WT; p = 0.03). While IL-1β was not significantly different, this is consistent with known challenges in detecting circulating IL-1β in IL-1-driven conditions^56^. Following GMA, IL-1α and other markers of IL-1 pathway activity were reduced to WT levels. IL-1α and IL-1β signal through the same receptor and activate overlapping inflammatory programs; IL-1α can act as an upstream alarmin that primes IL-1β responses, which are transient and often not systemically detectable^56,57^. Thus, reduced IL-1α after GMA indicated attenuation of IL-1-driven inflammation. Clinically, IL-1 blockade (anakinra or canakinumab) appears to reduce flare frequency and symptom burden in FOP patients; however, whether IL-1 pathway inhibition limits the formation of new HO has not been directly tested^58,59^. Furthermore, our prior findings suggest that suppression of IL-1-mediated inflammation may underlie the protective effects of GMA. We directly tested whether antibody-mediated IL-1β blockade using a murine anti-IL-1β antibody (01BSUR) analogous to canakinumab (human anti-IL-1β) was sufficient to reduce trauma-induced HO in our FOP mouse model. Because spontaneous HO is variable and unpredictable, and repeated manipulation can induce HO in the FOP mice, we used a standardized trauma model to test the effects of IL-1β inhibition.

FOP mice were subjected to a localized blunt-force injury in which a 60-g weight was dropped from a height of 20 cm onto the gastrocnemius muscle^60^, delivering approximately 16 N of force over a ∼10 mm² area (**Figure 4A**). This standardized injury better recapitulates the types of soft-tissue injuries often experienced by patients with FOP (i.e. after a fall), without the need of BMP augmentation or the use of local toxins^60,61^.

**Figure 4:**
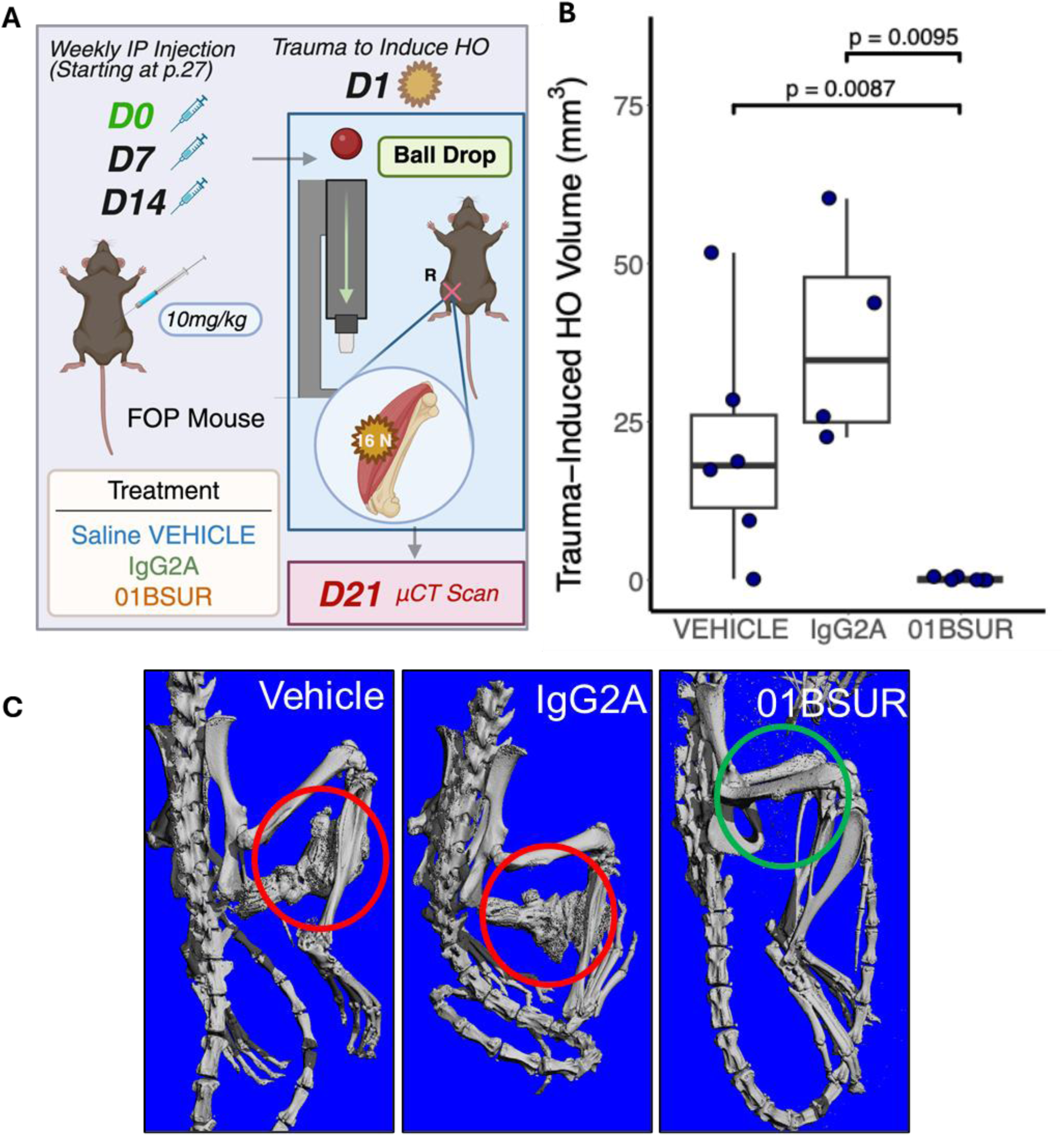
Pharmacologic Blockade of IL-1 Prevents Trauma-Induced HO. **(A)** Experimental design of the anti-IL-1β antibody study in the FOP mouse model. Mice received weekly intraperitoneal injections beginning at P27 (day 0) of either vehicle (saline), IgG2A isotype control antibody, or the anti-IL-1β monoclonal antibody 01BSUR (10 mg/kg). Trauma-induced HO was generated using a ball-drop injury model applied to the right hind limb at P28. HO formation was quantified by µCT at day 21. Created with Biorender.com. **(B)** Quantification of trauma-induced HO volume measured by µCT. Each point represents an individual mouse, and boxes show group distributions. Treatment with the anti-IL-1β antibody 01BSUR significantly reduced HO volume compared with both vehicle-treated mice (p = 0.0087) and IgG2A isotype controls (p = 0.0095). **(C)** Representative three-dimensional µCT reconstructions of skeletal structures from each treatment group. Areas of HO formation are highlighted. Extensive HO is observed in vehicle- and IgG2A-treated mice, whereas mice treated with the anti-IL-1β antibody show markedly reduced HO at the injury site. Representative images were chosen based on the mean HO volume of each group.

In control FOP mice, treatment with saline vehicle (VEH) or a non-specific IgG2A antibody resulted in substantial ectopic bone formation, with mean HO volumes of 20.99 mm³ and 38.05 mm³, respectively (**Figure 4B-C**). In contrast, FOP mice treated with the anti-IL-1 antibody 01BSUR exhibited a near-complete suppression of HO, with a mean HO volume of 0.199 mm³, corresponding to a 99.1% reduction relative to VEH (p = 0.009) and a 99.5% reduction relative to IgG2A (p = 0.01) (**Figure 4B-C**).

## DISCUSSION

Across chronic inflammatory diseases, gut microbiome dysbiosis has emerged as a driver of systemic immune activation through disrupted barrier integrity, altered metabolite signaling, and skewed innate immune responses^24–31^. FOP provides a unique opportunity to separate the impact of genetics and environmental factors, since FOP is a monogenic disorder driven by constitutive activation of *ACVR1* signaling. The clinical heterogeneity observed among FOP patients suggests that non-genetic modifiers play a critical role in shaping disease severity^36^. In this study, we provide evidence that the gut microbiome functions as an environmental regulator of systemic inflammation in FOP, modulating IL-1 pathway activity, HO burden, and ultimately survival.

Within our cohort, dietary patterns and GI symptoms did not differ between FOP patients and their cohabitating family members. The dietary similarity allowed us to do paired analyses of stool samples from siblings, revealing significant differences in overall gut microbial community composition. Patients with FOP exhibited higher-level taxonomic structure and species-level differences (**Figure 1D, Supplemental Table 3**), including depletion of taxa with reported immunoregulatory or anti-inflammatory associations and enrichment of those previously linked to inflammatory or pathobiont-associated behavior^37–46^ (**Supplemental Table 4**). In addition, the functional profiling of the gut microbiome further suggested enrichment of bacterial stress-response and metabolic pathways in FOP patients (**Supplemental Table 5**). Enrichment of ppGpp biosynthesis reflects activation of the bacterial stringent response (due to nutrient starvation, metabolic stress) which is associated with broad transcriptional reprogramming and changes in cell envelope and membrane-associated processes^47^. Increased prokaryotic TCA cycle pathways suggest more production of central metabolic intermediates, including succinate and fumarate, which are established as immunometabolic signaling molecules related to inflammation^48^. Additionally, enrichment of microbial arginine biosynthesis pathways suggests alterations in nitrogen metabolism that could influence host arginine availability, a key regulator of immune function, nitric oxide production, and inflammation^49^. Although these findings are limited by sample size and do not establish causality, the use of cohabitating sibling-matched pairs minimizes environmental factors and supports the idea that FOP is associated with altered gut microbial community structure, independent of overt gastrointestinal disease or dietary differences.

In the *PDGFRα-Cre;ACVR1^R206H^* mouse model^50^, baseline fecal microbial diversity and overall community composition did not differ significantly between FOP and littermate control (WT) mice (**Figure 2C**). Although species-level differential abundance analysis identified select taxa that differed between genotypes (**Supplemental Figure 2A-B**), Kegg-orthology (KO)-level data revealed only a small number of significantly altered microbial functions (**Supplemental Figure 2C-D**). These findings indicate that the restricted expression of *ACVR1^R206H^* in our mouse model (to PDGFRα expressing cells) is likely not sufficient to induce significant changes in gut microbial structure or function under controlled housing and dietary conditions. This raises the intriguing possibility that GI-tract activation of *ACVR1* may impact microbial composition, an area that has yet to be explored.

Despite no significant baseline microbiome differences, antibiotic-mediated GMA significantly reduced spontaneous HO volume in FOP mice (**Figure 2D-E**), suggesting that microbiome-derived signals are regulators of ectopic bone formation. The reduction in HO occurred despite no changes in plasma MMP-9 levels (**Figure 3A**), excluding a direct antibiotic effect on this previously implicated mediator^53^ and supporting an indirect mechanism by immune modulation.

Spontaneous and cumulative HO in this FOP mouse model is associated with early mortality, which was significantly improved by GMA-treatment (**Figure 2F**). Although the precise causes of mortality in this model are likely multifactorial and arise from the progressive HO accumulation, the survival benefit observed with GMA was associated with a reduction in systemic inflammation markers. At baseline, FOP mice exhibited elevated circulating levels of several pro-inflammatory and pro-fibrotic cytokines compared with WT controls, including IL-1α, IL-2, IL-11, and MIP-1α, (**Figure 3B**), consistent with prior reports in human FOP serum and iPSC-derived macrophages (**Supplemental Table** 7). Following GMA, many of these baseline differences, most notably in IL-1α, were attenuated (**Figure 3B-C**). In addition, within-genotype comparisons showed reductions in IL-1α, IL-6, eotaxin, and MIP-1β in FOP mice after GMA (**Figure 3B**). These findings indicate that the gut microbiome contributes to systemic inflammatory signaling in FOP and position the IL-1 pathway as a prominent downstream effector of microbiome-dependent immune activation.

Inflammatory flare-ups are a defining feature of FOP, often triggered by tissue injury and preceding HO formation^8^. The *ACVR1^R206H^*mutation enhances inflammatory responses^16,19,21,62,63^, and the IL-1 pathway has emerged as a central early mediator of HO across genetic, traumatic, and neurogenic contexts^58,59,64,65^. IL-1 signaling has been shown to promote HO by inducing RUNX2-mediated osteogenic differentiation in FAPs and is elevated early after HO-forming injury, supporting its role as both a driver and biomarker of HO^64,65^. Human iPSC-derived macrophages carrying *ACVR1^R206H^* exhibit prolonged M1-like inflammatory activation with elevated baseline IL-1α^19^. Furthermore, clinical case-series in patients with FOP show that IL-1 blockade with anakinra and/or canakinumab can reduce flare frequency and symptom burden^58,59^. One study with tofacitnib, a JAK inhibitor that also reduces IL-1 pathway activity, also demonstrated reduction in flare activity^66^. However, whether IL-1 blockade is associated with decreased HO formation in FOP patients remained unknown.

Our results showed that antibody-mediated IL-1β blockade nearly abolished trauma-induced HO in FOP mice, resulting in a greater than 99% reduction in HO volume (**Figure 4B-C**). The magnitude of this effect suggests that IL-1 pathway activity is a functional driver of injury-induced HO in FOP. These findings also suggest that the IL-1 pathway may integrate multiple upstream inflammatory inputs, including microbiome-derived signals and inflammation-related signals, to promote *ACVR1*-mediated osteogenic differentiation following injury.

Our study has several limitations. The human studies were limited by the small sample size, which is not unexpected due to the rarity of FOP. This limited our ability to identify if specific species may be drivers of FOP inflammation and HO in humans. In the FOP mouse studies, the severity of the phenotype significantly reduced survival, which limited the duration of experiments and precluded our ability to reliably assess long-term outcomes of GMA beyond the 4 weeks of treatment in our study design, and necessitated the use of adolescent mice. Additionally, while GMA was effective in reducing HO in mice, and manipulation of the gut microbiome and subsequent inflammatory modulation appear to be major contributors to this effect, potential non-microbial effects of the antibiotics beyond MMP-9^53^ remain unknown.

Although our results identify a link between the gut microbiome, IL-1 pathway activity, HO formation, and survival (**Figure 5**), whether GMA might have a benefit for reducing post-traumatic HO in FOP patients remains to be elucidated. GMA here was also employed as an experimental perturbation in a preclinical mouse model to interrogate the contribution of gut-derived signals to HO. The long-term use of broad-spectrum antibiotics for the prevention or treatment of HO in patients with FOP or other forms of HO is not currently warranted, given the known risks of antibiotic resistance and gut dysbiosis. However, these findings suggest that modulation of the gut microbiome through alternative, clinically relevant strategies, such as targeted microbial interventions, dietary modulation, pre- and probiotic approaches, or even a brief course of broad-spectrum antibiotics in the setting of trauma, may represent useful therapeutic strategies to be studied for influencing inflammation and subsequently HO.

**Figure 5.**
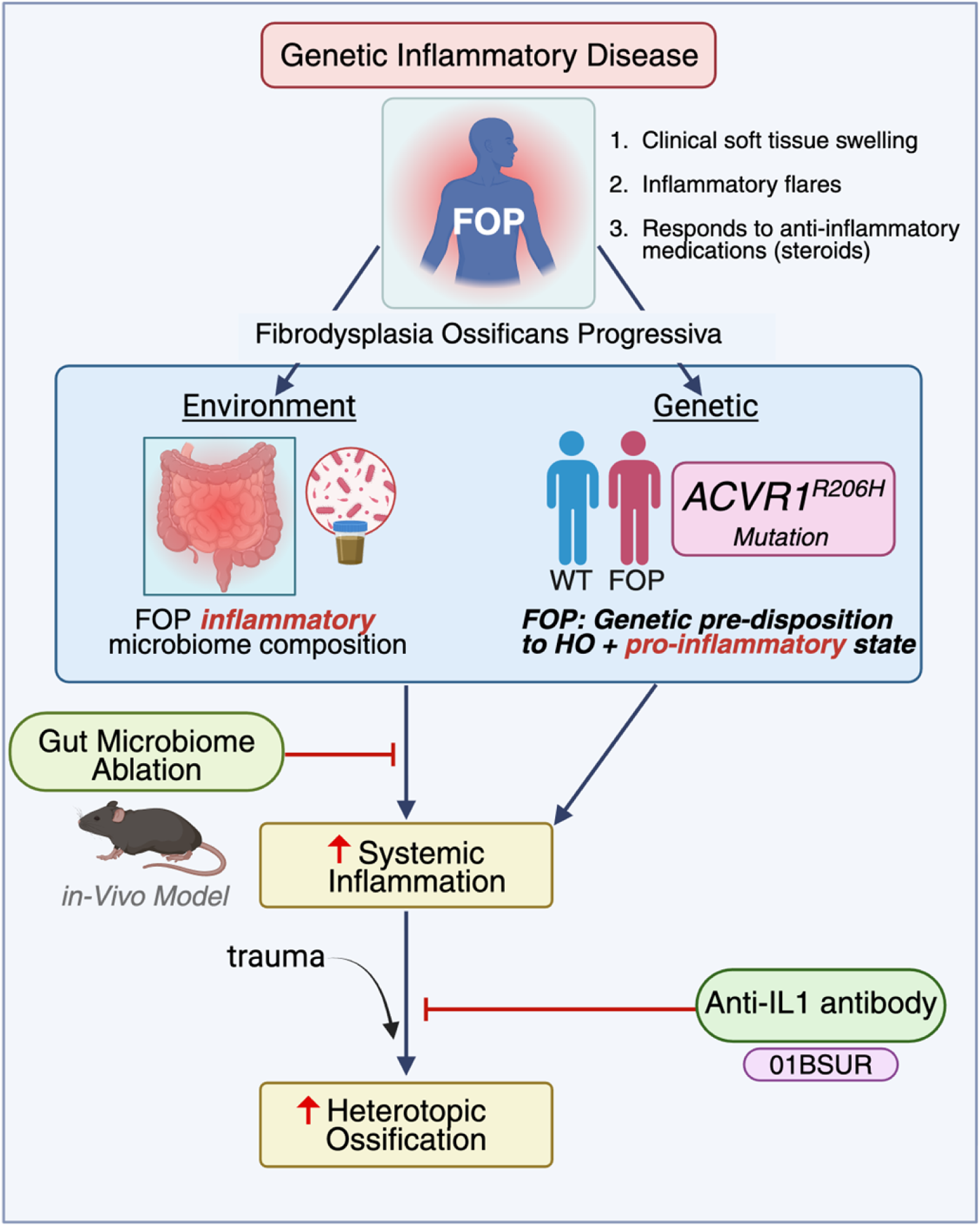
Proposed model linking the gut microbiome, inflammation, and HO in FOP. FOP is a genetic inflammatory disease caused by activating mutations in *ACVR1* (R206H), resulting in a pro-inflammatory state and predisposition to HO. Environmental factors, including the gut microbiome, further shape systemic inflammation, with FOP-associated microbial communities promoting inflammatory signaling. Gut microbiome ablation (GMA) *in vivo* reduces systemic inflammation (IL-1), highlighting the contribution of microbiota-derived signals. Following trauma, elevated inflammatory signaling drives ectopic bone formation. Targeting inflammatory pathways, including IL-1 signaling with neutralizing antibodies (e.g., 01BSUR), mitigates HO development. This model integrates genetic and environmental contributions to inflammation-driven HO in FOP. Created with Biorender.com.

Notably, many of the inflammatory features found in FOP are likely relevant to other forms of non-genetic HO^20^. Attempts to identify genetic drivers of non-genetic HO have been largely unsuccessful, as multiple genes are potentially implicated^67–69^. The observations of IL-1 pathway relevance in neurogenic, traumatic, genetic, and clinical FOP contexts suggest that this pathway is likely a major driver of HO initiation and propagation^58,59,64,65^, and thus an important clinical target. Future studies are also needed to identify the specific microbiome-derived signals that link to the IL-1 pathway. Furthermore, larger scale longitudinal human studies incorporating repeated stool sampling, flare tracking, and treatment data will be valuable to determine whether microbiome features may be a useful marker to predict disease activity or therapeutic responses in patients with FOP or with other forms of HO.

In summary, these data identify the gut microbiome and IL-1 pathways as modifiable drivers of HO and suggest that targeting microbiome-immune interactions may represent a therapeutic strategy for FOP and other inflammation-driven disorders of ectopic ossification

## MATERIALS AND METHODS

### Human Study Cohort

The human study was approved by the UCSF Internal Review Board. Written informed consent was obtained from all participants and/or legal guardians. Participants with FOP were enlisted in this study between June 2017 and December 2018 (**Figure 1A**). All patients met clinical diagnostic criteria for classical FOP, including classical skeletal hallmarks, a history of FOP flares, and congenital malformations of the great toes (hallux valgus). In addition, healthy family controls were included in the study. These individuals were biological siblings of patients and lived in the same household, except for one control, who was a spouse living in the same household. For the subset of participants for the stool analyses, candidates were excluded if they did not have a matching sibling; had any current infection or antibiotic use within one month before participating in the study; or were taking an investigational or known disease-modifying drug (e.g., palovarotene). Brief clinical summaries are provided in **Supplemental Table 8 and Figure 1A**.

### Survey Collection and Analysis

Consenting subjects were surveyed to obtain detailed information about food preferences, eating habits, and GI symptoms using questionnaires adapted from published validated surveys such as the GI-PROMISE, ^70–72^. The questionnaires were provided in an electronic format through REDCap, a secure survey and database web application ^73^. Cumulative joint involvement scores (CAJIS) were collected on FOP patients via video visits or in person with study staff to assess the functional status of FOP disease progression ^36^. One subject was unable to complete the CAJIS assessment within the expected timeframe. Data were analyzed using Stata Statistics (College Station, TX, version 16). Survey responses were dichotomized and differences in frequency patterns of categorical variables (eg, GI symptoms, food preferences) were examined with Fisher’s exact tests (p < 0.05 considered significant).

### Human Stool Sample Collection

For the microbiome studies, stool was collected from patients with FOP who did not have a clinically active flare at the time of stool collection.

Participants were provided with a kit for home fecal sample collection (Zymo Research DNA/RNA Shield Fecal Collection Tube, Catalog No. R1101). Collected specimens were frozen at -20°C by the participant, shipped overnight to University of California, San Francisco, and frozen at -80°C on arrival for future processing. Total DNA (microbial and host) was extracted using the Zymogen Quick DNA Fecal/Soil Microbe Miniprep Kit (Catalog No. D6010). Fecal samples were considered unusable if inappropriate sample preservation was documented. Samples that yielded low concentrations of microbial DNA were extracted a second time. If the low DNA yield persisted, these samples were also excluded from further analysis.

Twelve participants did not complete the full survey and sample collection (due to loss of interest in the study, failure to return forms in the expected timeframe, unwillingness to collect the stool sample, or failed stool sample collection/processing); these were excluded from the microbiome analyses (**Figure 1A**).

### Human Stool Metagenomic Sequencing and Assembly

To minimize environmental variation in human microbiome analyses, only paired samples consisting of individuals with FOP and their unaffected sibling controls were analyzed (n = 7 pairs; 14 individuals total). Fecal DNA was extracted using a bead-beating protocol with the Fecal DNA Miniprep Kit (D6010, Zymo Research) according to the manufacturer’s instructions. Sequencing libraries were prepared by the UCSF Institute for Human Genetics (IHG) Core using TruSeq adaptors and sequenced on an Illumina HiSeq 4000 or NovaSeq 6000 platform to generate 150-bp paired-end reads with a target depth of approximately 50 million reads per sample. Shotgun metagenomic sequencing produced consistently high read depth across all samples (approximately 65-140 million reads per sample) with no major outliers, indicating sufficient and balanced sequencing coverage for downstream taxonomic and functional analyses (**Supplemental Figure 5**).

Taxonomic profiling was performed using the CosmosID bioinformatics platform (CosmosID, Germantown, MD, USA) and its proprietary analytical tools. The CosmosID taxonomic profiling algorithm consists of two main phases: a pre-computation phase for building a curated microbial biomarker database and a computational phase for classifying sequencing reads against this database.

During the pre-computation phase, a curated collection of high-quality microbial genomes is assembled from publicly available databases, primarily NCBI RefSeq, with consideration of genome completeness, contamination, and assembly quality. The resulting reference database contains more than 25,000 microbial species spanning multiple kingdoms, including Bacteria, Viruses, Fungi, Protists, and Phages. Genomes in this database are decomposed into variable-length n-mers, which are then categorized as either shared or unique genomic biomarkers. These biomarkers are organized within a phylogenetic tree-like data structure in which internal nodes represent shared biomarkers across taxa and terminal leaves represent unique genomic signatures for individual microbial genomes.

In the per-sample computational phase, millions of sequencing reads are queried against this phylogenetic biomarker database. Reads are first decomposed into k-mer sets that are matched against the biomarker tree. Exact matches between query k-mers and reference biomarkers are then identified to determine taxonomic assignment. Taxonomic classification and relative abundance estimates are calculated using composite statistics derived from biomarker matches, coverage depth, and genome size, which are integrated into a proprietary abundance scoring system. The core algorithm underlying the CosmosID platform is protected under US and European patents (US10108778B2, US20200294628A1, and ES2899879T3).

### Stool Statistical Analysis

Differential abundance analysis was performed using the DESeq2 algorithm (version 1.24.0) with default parameters, including median-of-ratios normalization, negative binomial generalized linear modeling, and Wald tests for significance, to identify differences in the relative abundance of microbial taxa and Kyoto Encyclopedia of Genes and Genomes (KEGG) functional features, including KEGG orthologs (KOs), modules, and pathways, between responders and non-responders in the pre-treatment microbiome. P-values were adjusted for multiple hypothesis testing using the Benjamini-Hochberg false discovery rate (FDR) procedure.

Prior to differential abundance testing, features with minimal signal were excluded from analysis; specifically, features were required to be present in at least three samples from at least one study group with an abundance greater than one-fifth of the average signal across all features. Overall differences in gut microbiome composition were assessed using Bray–Curtis distance-based permutational multivariate analysis of variance (PERMANOVA) based on the relative abundance of microbial taxa and KEGG orthologs.

Linear Discriminant Analysis Effect Size figures were generated using LEfSe ^74^, based on taxonomic matrices from CosmosID and annotations from MetaCyc ^75^ and GO ^76^. LEfSe is calculated with a Kruskal-Wallis alpha value of 0.05, a Wilcoxon alpha value of 0.05, and a logarithmic LDA score threshold of 2.0. In the LEfSe figures, red bars to the right convey that the feature in that group is more abundant in the “red” group than the other. Blue bars to the left convey that the organism is more abundant in the “blue” group.

Stacked bar figures were generated using taxonomic matrices from CosmosID and annotations from MetaCyc ^75^ and GO ^76^. Stacked bar figures for each group were generated using the R packages ggplot2 and ggpubr. Heatmap figures were generated using taxonomic matrices from CosmosID and annotations from MetaCyc ^75^ and GO ^76^ using the heatmaply R package. Alpha diversity boxplots were calculated from the taxonomic matrices from CosmosID and annotations from MetaCyc ^75^ and GO ^76^. Simpson alpha diversity was calculated in R using the R package Vegan ^77^. Wilcoxon Rank-Sum tests were performed between groups using the R package ggsignif. Boxplots with overlaid significance in p-value format were generated using the R package ggplot2.

Beta Diversity Principal Coordinate Analyses were calculated from the taxonomic matrices from CosmosID and annotations from MetaCyc ^75^ and GO ^76^. Bray-Curtis diversity was calculated in R using the R package Vegan ^77^ with the functions vegdist, and PCoA tables were generated using ape’s function pcoa. PERMANOVA tests for each distance matrix were generated using Vegan’s ^77^ function adonis2, and beta dispersion was calculated on the Bray-Curtis dissimilarities using the betadisper function. Plots were visualized using the R package ggpubr, and comparisons were conducted for taxonomic analyses in both paired-sample and independent models.

### Analyses Generated

LEfSe Differential Abundance Analysis: Differential feature and organism abundance between cohorts is calculated using three methods: Kruskal-Wallis sum-rank test, Wilcoxon rank-sum test, and Linear Discriminant Analysis. Analysis was performed at all taxonomic levels as well as for GO ^76^ and MetaCyc ^75^ functional analyses. Relative abundance taxonomic matrices and counts-per-million functional matrices were used as input. All features shown meet p≤0.05 for Kruskal-Wallis and Wilcoxon tests and have an LDAscorewhere effect-size ≥2.0.

Relative Abundance Stacked Bar and Heatmaps Charts: Stacked bar figures show the top organisms at each taxonomic level. Analysis was performed at all taxonomic levels as well as for GO ^76^ and MetaCyc ^75^ functional analyses. Relative abundance taxonomic matrices and counts-per-million functional matrices were used as input.

Alpha Diversity Boxplots with Wilcoxon Rank-Sum Test: Shannon and Simpson diversities indicate richness as well as the evenness of the organisms within a sample, where a higher value indicates both higher species evenness and richness (Simpson diversities give lower weighting to rarer species, while Shannon is considered “unbiased”). Shannon entropy is the natural logarithm of Shannon diversity and is commonly used to describe community composition. Boxplot figures were generated at all taxonomic levels. Relative abundance taxonomic matrices and counts- per-million functional matrices were used as input. Each of the cohort pairs being analyzed were compared using the Wilcoxon Rank-Sum Test.

Beta Diversity PCoA with PERMANOVA: Principal Coordinates Analysis (PCoA) shows the difference between entire microbiomes, with each point representing the community of a single sample. Greater distance between points indicates more dissimilar communities. Bray-Curtis dissimilarity accounts for the presence or abundance of each organism in addition to their relative abundances. PERMANOVA test results were included for each cohort pair. PCoAs were generated at all taxonomic levels and for MetaCyc ^75^ and GO ^76^ functional analyses.

### Antibiotic Mouse Studies – Study Cohort

All mouse studies were approved by the UCSF Institutional Animal Care and Use Committee (IACUC) and Laboratory Animal Research Center (LARC) at the University of California, San Francisco*. PDGFRα-cre/ACVR1^R206H^* (abbreviated as FOP) mice ^13,50^ were weaned at 21 days of age from parents that received normal water (without antibiotics). A total of 24 mice in 4 cohorts were used: 6 FOP mice weaned onto antibiotics, 6 FOP mice weaned without antibiotics, 6 control mice weaned onto antibiotics, and 6 control mice weaned without antibiotics. The control animals included *PDGFRα-cre* and *ACVR1^R206H^* single transgenics, as well as wildtype littermates (WT), since there were no significant phenotypic differences among these control genotypes. All littermates were housed together and separated only by sex after weaning.

The broad-spectrum antibiotic cocktail consisted of 1 g/L ampicillin, 0.5 g/L metronidazole, 0.5 g/L neomycin, and 0.5 g/L vancomycin dissolved in 1 L of autoclaved deionized water. This antibiotic water (ABX) was changed twice per week.

### FOP Mouse Kaplan-Meier Survival Curve

FOP mice were administered the broad-spectrum ABX cocktail beginning at postnatal day 21 (weaning) to achieve gut microbiome ablation (GMA). Untreated FOP littermates served as controls. Survival was recorded daily until natural death, or if the UCSF veterinarian recommended euthanasia for meeting endpoint criteria. All mice were included in the analysis.

Survival analyses were performed in R using the survival, survminer, and ggplot2 packages. Survival time was defined as the number of days each animal remained alive, and death was treated as the event for all mice. A survival object was constructed using survival time and event status, and mice were categorized into four groups based on sex and treatment (male FOP, male FOP GMA, female FOP, female FOP GMA). Kaplan-Meier survival curves were generated for each group, and visualizations were produced using survminer. Log-rank tests were used to compare survival distributions between FOP and FOP GMA groups. Test statistics were derived from chi-square distributions with one degree of freedom, and two-sided p-values were reported without adjustment for multiple comparisons.

### Mouse microCT Imaging and Analysis

Full-body microCT scans were done on mice at 4 weeks post-GMA. Skeletons were collected in 50mL conical tubes and preserved in formalin for 24 hours at 4 °C and then changed to 70% ethanol solution and shipped overnight with dry ice via FedEx to Emory University in Atlanta, Georgia, USA. Both 3D mouse imaging and analysis were done on the microCT 50 SCANCO Medical AG scanner and associated software (Brüttisellen, Switzerland). Images were obtained at 50 kV, 200 µA, and 12.4 µm isotropic voxel spacing, with a 450 m/sec integration time and 1000 projections through a 35 mm FOV. The manufacturer’s 1200 mgHA/ccm beam hardening correction was applied during image reconstruction.

To specifically select heterotopic ossification bone on the scans, manual outlines contours were created around the desired areas. Vertebral HO distal to the pelvic girdle was not analyzed. A threshold of 250 mgHA/ccm with noise filter settings of Support 1 and Sigma 0.8 was then used to quantify the mineralized volume. Quantification of the resulting 3D structure was calculated by direct voxel counting. The mean tissue density was calculated by dividing the total bone mineral content by the bone volume (BMC/BV). Total mineralized density (TMD) (mg HA/ccm) and mineralized HO (mm^3^) were additionally quantified for both FOP mice treated with ABX and untreated FOP mice. When visualizing the microCT figure, each HO image was overlaid or superimposed on the segmented image of the whole-body skeleton. All µCT images were evaluated individually to identify HO presence in axial and appendicular regions. The appendicular region included HO in the limbs and extremities. The axial region was subdivided into mandibular HO at the jaw and upper spinal HO below the skull. HO presence in each region was recorded as a binary outcome (yes/no).

All analyses were performed in R using the packages readxl, dplyr, ggplot2, and ggpubr. HO volume data were imported from an Excel spreadsheet. One FOP outlier, mouse ID 789, was excluded from the analysis because the animal did not develop any HO, consistent with a likely genotyping error. After filtering, HO volumes were compared between FOP and FOP GMA groups. Normality was assessed within each group using the Shapiro-Wilk test, and equality of variances was evaluated using an F-test. Both groups passed normality testing; however, because variances were unequal between groups, a two-sided Welch’s t-test was used to compare HO volume between FOP and FOP GMA mice. Summary statistics are reported as mean ± SD, and percent reduction in HO volume with GMA was calculated relative to the FOP group mean.

### Mouse Stool Sample Collection

Upon weaning, at least two stool pellets per mouse were collected into 1mL Eppendorf tubes and stored at –80 °C. Samples were collected in the morning. These samples served as baseline measures which were recorded as Timepoint 1 (t1). Four weeks post-wean (P49), mice were sacrificed and harvested, which was recorded as Timepoint 2 (t2). Two stool pellets were collected immediately before euthanasia and harvest, and blood was collected via cardiac puncture and centrifuged at 10000 RPM for 30 minutes to isolate plasma for cytokine quantification and analysis regarding inflammation. Carcasses were analyzed by micro X-ray computed tomography (microCT) for HO quantification (details below). Stool and plasma samples were collected in 1 mL Eppendorf tubes and stored at -80 °C.

### Stool DNA Extraction and Library Preparation

Fecal samples from mice (n = 48), representing six mice from four experimental groups at two different timepoints (baseline/pre-treatment and 4 weeks post-GMA), were collected for DNA extraction and subsequent shotgun metagenome sequencing. DNA extraction was performed using the CTAB (Cetyltrimethylammonium bromide) extraction protocol optimized at the Microbial Genomics CoLab Plug-in of the Benioff Center for Microbiome Medicine (BCMM) at UCSF as detailed in a published article ^78^. The extracted DNA was quantified using a broad range Qubit dsDNA Quantification Assay Kits (Thermofisher Scientific, USA). Shotgun metagenomic libraries were prepared using the Illumina DNA Prep kit (Catalog # 20060059, Illumina, San Diego, CA, USA) employing unique barcode sequences. The final libraries were pooled at equimolar concentrations and analyzed on a Bioanalyzer for quality prior to metagenomic sequencing at the UCSF Center for Advanced Technology (CAT) facility. The pooled libraries were sequenced using the Illumina NovaSeq 6000 in a 2×150 bp paired-end run protocol.

### Shotgun Metagenome Sequencing and Data Processing

#### Sequencing

Shotgun metagenomic sequencing data showed consistently high base-call quality across reads (**Supplemental Figure 6A-B**). FASTQC mean Phred scores remained largely above Q30 across the full read length, with only a modest decline toward the final cycles (**Supplemental Figure 6A**), and per-sequence quality scores were tightly distributed in the high-quality range (**Supplemental Figure 6B**). Raw FASTQ files were used for all downstream analyses. Kneaddata processing reduced read counts across stages as expected (input → trimmed → final), with substantial numbers of reads retained per sample after trimming and final filtering (**Supplemental Figure 6C**).

#### Computing Environment and Pipeline

All analyses were performed on the UCSF Wynton High-Performance Computing (HPC) cluster. All processing steps (quality control, host removal, taxonomic classification, functional profiling) were executed on Wynton compute nodes.

#### Quality Control and Preprocessing

Initial quality assessment of raw FASTQ files was performed using FastQC (v0.12.1). Reads were then trimmed and filtered with KneadData (v0.12.3), which uses Trimmomatic (v0.39) for adapter removal and quality trimming. Quality filtering was performed using a sliding 4-bp window, and reads were trimmed when the average Phred score within the window dropped below 20. Reads shorter than 50 bp after trimming, as well as those containing residual adapter sequence, were removed. KneadData also incorporates Tandem Repeat Finder (TRF), which was used to remove over-represented low-complexity sequences prior to downstream taxonomic profiling.

#### Host DNA Removal

To remove host-derived sequences, KneadData aligned all reads to the C57BL/6NJ mouse reference genome using Bowtie2 (v2.5.1). Host-mapped reads were removed, and only unmapped, high-quality reads were retained for downstream taxonomic and functional profiling. Bowtie2 was run using the very-sensitive-local preset with dovetail alignments enabled to maximize host read detection.

#### Taxonomic Profiling

Taxonomic classification was performed using Kraken2 (v2.1.3) with the standard Kraken2 database (January 4th, 2023 build). Kraken2 uses exact k-mer matching to assign reads to taxa across bacterial, archaeal, viral, and eukaryotic lineages. Species-level abundance estimates were refined using Bracken (v2.9), which applies Bayesian k-mer redistribution to improve quantitative accuracy. Kraken2 report files and Bracken abundance tables were used to generate species-level count matrices for downstream analysis. Taxa present in <10% of samples or with extremely low estimated abundance were removed prior to downstream analysis. For validation, a complementary marker-gene-based taxonomic profile was also generated using MetaPhlAn 4.0.6 using the mpa_vOct22_CHOCOPhlAnSGB_202403 Bowtie2 database. Kraken2/Bracken outputs were used as the primary taxonomic dataset. Relative abundance QC plots are shown in **Supplemental Figure 7**.

#### Functional Profiling

Functional profiling was performed using HUMAnN 3.8, which identifies microbial gene families and metabolic pathways through a tiered alignment strategy:

1. Nucleotide alignment of reads to the ChocoPhlAn pangenome database (full_chocophlan.v201901_v31.tar.gz).
2. Translated search of unmapped reads against the UniRef90 protein database (uniref90_annotated_v201901b_full.tar.gz), enabling functional detection independent of reference genomes.
3. Pathway reconstruction using MetaCyc annotations.

Gene families were regrouped into KEGG Orthologs (KOs) using HUMAnN’s humann_regroup_table utility, and all functional abundance tables were normalized to copies per million (CPM). KO groups detected in <10% of samples were excluded. The final functional dataset included 2771 of KO groups.

### Downstream and Statistical Analyses

#### Alpha diversity analysis

Alpha diversity was quantified at the species level using size-factor-normalized count data in R. Richness was calculated as the number of observed species using the vegan package, and within-sample diversity was estimated using the Shannon diversity index. Group differences in richness and Shannon diversity were evaluated using two-sided Wilcoxon rank-sum tests, and unadjusted p-values were reported.

To assess overall effects of genotype and GMA while accounting for repeated measurements within mice, linear mixed-effects models were fit for each alpha diversity metric using lme4, with experimental group specified as a fixed effect and MouseID included as a random intercept. Model significance was evaluated using F-tests derived from the corresponding ANOVA. Alpha diversity distributions were visualized using ggplot2 with jittered points overlaid on boxplots.

#### Beta diversity analysis

Beta diversity was assessed using Bray-Curtis dissimilarity calculated from species-level normalized abundance tables. Principal Coordinates Analysis (PCoA) was performed using the phyloseq package to visualize community structure, and the first two axes were used for plotting with ggplot2. For the GMA comparison, analyses were restricted to WT_GMA and FOP_GMA (Abx) samples, and one extreme outlier (FOP_Abx_11) was excluded prior to analysis. Differences in community composition were tested using PERMANOVA implemented in adonis2 from the vegan package, modeling Group × Sex and Timepoint as predictors. Baseline (non-GMA) differences between WT and FOP mice were evaluated separately using Bray-Curtis PCoA, followed by PERMANOVA models testing genotype and sex effects, including WT vs FOP overall, FOP males vs females, WT males vs females, and sex-matched WT vs FOP contrasts. All PERMANOVA tests used 999 permutations.

#### Differential abundance testing

Differential abundance analyses were conducted in R using phyloseq, DESeq2, dplyr, ggplot2, ggprism, pheatmap, and scales. Taxonomic count data were normalized using DESeq2 size factors (sfType = “poscounts”) and agglomerated to Kingdom, Phylum, or Class levels for visualization. Relative abundances were calculated on a per-sample basis and used to generate stacked bar plots stratified by experimental group.

Differential abundance testing was performed with DESeq2 on taxa present in more than 30% of samples. Models were constructed to evaluate effects of genotype (FOP vs. WT), antibiotic treatment (GMA vs. non-GMA), and timepoint. Wald tests were used to estimate log₂ fold changes, and p-values were adjusted using the Benjamini-Hochberg procedure, with padj < 0.05 considered statistically significant.

Volcano plots were generated using ggplot2 and ggprism, displaying log₂ fold change versus – log₁₀(adjusted p-value). Significant differentially abundant taxa were assigned to species using the phyloseq taxonomy table, aggregated to species-level counts, log₁₀-transformed, and standardized by row-wise z-scoring. Heatmaps were generated using pheatmap with Euclidean distance and complete-linkage clustering, with phylum-level annotations included.

#### Functional Profiling and Differential Abundance Analysis (KEGG KOs)

Functional profiling was performed using HUMAnN 3.0 ^79^. Gene family abundances were regrouped to KEGG Orthologs (KOs) and normalized to copies per million (CPM). KOs containing composite identifiers or non-informative labels (e.g., UNMAPPED, UNGROUPED) were removed. Samples with fewer than 10% detected KOs and KOs present in fewer than 10% of samples were excluded prior to analysis.

KO abundance tables and KEGG annotations were imported into R for downstream analyses using phyloseq (1.44.0) ^80^, DESeq2 (1.40.2) ^81^, pheatmap, ggplot2, and ggprism. Differential abundance between FOP and WT groups was evaluated using DESeq2, with size factors estimated using the poscounts method and a model testing KO-level differences between groups. Multiple testing correction was applied using the Benjamini-Hochberg false discovery rate (FDR) procedure, and KOs with FDR-adjusted p-values < 0.05 were considered significant.

Volcano plots were generated using ggplot2 and ggprism, displaying log₂ fold change versus -log₁₀(FDR-adjusted p-value) for individual KOs. Significant KOs were highlighted and colored by KO subclass for visualization.

Heatmaps were constructed from the set of significantly different KOs. DESeq2-normalized KO abundances were averaged within each experimental group, log₁₀-transformed, and standardized by row z-scoring to scale each KO relative to its own mean and variance across conditions. The resulting z-score matrix was visualized using pheatmap, with hierarchical clustering of rows (Euclidean distance, complete linkage) and annotation of columns by experimental group.

### Cytokine Studies

Mice were anesthetized and blood was collected by cardiac puncture using a sterile needle pre-coated with heparin. 0.5-1ml of blood was slowly and steadily withdrawn to prevent vessel collapse. Following blood collection, the animal was euthanized, and the collected blood was immediately transferred to ice. The collected blood samples were then centrifuged at 2,000 g for 10 minutes at 40C to separate plasma and cells. Plasma was then aliquoted and stored at -80C until analysis. For Cytokine analysis, samples were sent to Eve Technologies (Canada) for Luminex analysis with Mouse Cytokine 44-Plex Discovery Assay®.

Cytokine measurements were imported into R from Excel and organized by experimental group (WT, WT GMA, FOP, FOP GMA). Heatmaps were generated using the ComplexHeatmap package in R. Group means were calculated for each cytokine. To assess differences between conditions, log₂ fold-change values were derived either by comparing FOP GMA directly to FOP (single-contrast analysis) or by normalizing all groups to the WT group mean (multi-group analysis). To avoid division by zero, a small constant (pseudocount; 1×10^-6^) was added to all values prior to fold-change calculation.

### MMP-9 ELISA

FOP and WT mice were administered a broad-spectrum antibiotic cocktail consisting of ampicillin (1 g/L), neomycin (0.5 g/L), metronidazole (0.5 g/L), and vancomycin (0.5 g/L) in their drinking water for one week. Antibiotic-containing water was replaced every three days. Following treatment, mice were anesthetized with isoflurane, and blood was collected via cardiac puncture prior to euthanasia by exsanguination followed by cervical dislocation. Whole blood was incubated at room temperature for 2 hours to allow clot formation, then centrifuged at 2,000 × g for 20 minutes. Plasma was separated, aliquoted, and stored until analysis. Plasma MMP-9 levels were quantified using the Quantikine Mouse Total MMP-9 Immunoassay (Fisher Scientific; catalog no. MMPT90), according to the manufacturer’s instructions.

The effects of genotype and condition on MMP-9 levels was determined using linear mixed-effects model using the lmer function from the lmerTest package in R (v3.1.3). The model included fixed effects for genotype (control or FOP), condition (control or GMA), and their interaction (genotype × condition). Statistical significance was defined as p < 0.05. Analyses were conducted using R Version 2025.09.1+401.

### IL-1 STUDY

#### Animals and Genotyping

Littermates were genotyped at postnatal day 21 (P21). Mice carrying the FOP allele were identified, ear-tagged, and randomly assigned to treatment groups. Randomization was performed prior to initiation of dosing to generate a treatment matrix for all experimental conditions. IgG2A isotype antibody from R&D Systems (catalog no. MAB0031) was used as a control. The experimental antibody 01BSUR (monoclonal anti-IL-1β antibody, against mouse IL-1β, kindly provided by Novartis Pharma, AG) was included as an additional treatment condition. All antibodies were administered at a dose of 10 mg/kg (100 ng/µL) by intraperitoneal injection. Vehicle-treated animals received an equivalent injection volume of sterile saline. A total of 16 mice were used across the experimental groups: 01BSUR (n = 6), IgG2A (n = 4), and Vehicle (n = 6). The 01BSUR group consisted of 5 males and 1 female. The IgG2A group consisted of 1 male and 3 females. The Vehicle group contained 3 males and 3 females.

#### Dosing Schedule

At P26, all mice were weighed, and individual dosing volumes were calculated for administration on P27. Antibody and vehicle treatments were administered once weekly from P27 (day 0, D0) until the experimental endpoint. Mice were reweighed the day prior to each subsequent injection (P33 and P40), and dosing volumes were adjusted accordingly.

#### Ball Drop Injury Model

A ball drop-based noninvasive blunt injury method was used to elicit localized heterotopic ossification (HO). On P28 (day 1, D1), mice were anesthetized and positioned so that the right hind limb rested under a fixed pin. A weighted ball was released from a defined height onto the pin to deliver a controlled force to the limb. This procedure induces directed soft-tissue injury that promotes site-specific HO formation ^60^ (**Figure 4A**).

#### Animal Monitoring and Husbandry

Mice were monitored regularly throughout the study for health status, body condition, and treatment-related effects. Environmental enrichment, including oatmeal bowls, water gel, and foraging mix, was provided as needed. Any animals that died before the planned harvest date were collected and stored at −20°C for subsequent analysis. *In Vivo* collection and postmortem processing for micro-computed tomography (µCT) at P49, mice were euthanized and processed for blood and skeletal tissue collection. Prior to harvest, all dissection tools, collection tubes, and labels were prepared and sterilized. Mice were euthanized individually by CO₂ asphyxiation, and death was confirmed by the absence of a paw pinch reflex. Whole blood was collected immediately by cardiac puncture using a heparinized syringe and transferred to pre-labeled microcentrifuge tubes. Blood samples were kept on ice for approximately 20 minutes and subsequently centrifuged at 8000 rpm for 10 minutes to isolate plasma. Plasma aliquots were stored at –80 °C until analysis.

Following blood collection, carcasses were transferred for dissection and preparation for µCT imaging. After verifying mouse identification via ear tag, the ventral surface was wetted with 70% ethanol and incised longitudinally and transversely to expose the abdominal cavity. Internal organs, including abdominal viscera, diaphragm, heart, lungs, and thymus, were removed. The remaining carcass was carefully de-furred to preserve skeletal integrity and any heterotopic ossification present. The carcasses were placed per labeled storage container, submerged in 70% ethanol, and stored at 4 °C until µCT scanning.

#### MicroCT Imaging and Analysis

Both 3D mouse imaging and analysis were performed using the SCANCO Medical µCT50 scanner (SCANCO Medical AG, Brüttisellen, Switzerland) and the associated analysis software at UCSF. Images were acquired at 55 kV, 109 μA, with an isotropic voxel size of 20.0 μm, using an integration time of 500 ms and 1000 projections over a 35.28 mm field of view (FOV). Regions containing HO were acquired by manually tracing contours around the relevant anatomical structures. Mineralized tissue was segmented using a 173 mg HA/ccm threshold with a noise filter set to Support = 1 and Sigma = 0.8. Mineralized volume was obtained by direct voxel-based quantification of the resulting 3D reconstruction. Mean tissue density was determined by dividing the total bone mineral content by the corresponding bone volume (BMC/BV). For all FOP mice, total mineralized density (TMD; mg HA/ccm) and the volume of mineralized HO (mm³) were also measured. Quantifications were separated by experimental groups, and statistical significance between groups was determined by two-tailed T-tests.

## Acknowledgements

The authors thank Apsara Ram, Anastasia Diolintzi, Li Zhang, Rheinallt Jones, and Darren Dumlao, for their valuable technical discussions.

## Contributions

Project conception: HH, CF, LL, DP, ECH;

Funding: ECH, HH, DP;

Experimental execution: HH, CF, LL, KJ, SWT, KY, FVL, NH, AS;

Data analysis: AZ, CLG, HH, ED, TM, MH, SL, IM, SL, NH, KSP, CJH, DP, AS, ECH;

Drafting of the manuscript: HH, CF, LL, KJ, AZ, SWT, NH, ECH;

Editing of the manuscript: HH, KJ, DP, CJH, ECH, KSP;

Approval of final manuscript: all authors.

## Generative AI statement

Grammarly was used as a tool for editing and grammar suggestions in this manuscript. No other generative AI tools were used in the writing of this manuscript.

## Data and materials availability

Sequencing data generated during this study will be publicly available in the NCBI Sequence Read Archive (accession pending), including raw sequencing reads and associated metadata identifiers. The *ACVR1^R206H^* mouse was obtained from Dr. David Goldhamer’s laboratory under a materials transfer agreement (MTA) that does not allow redistribution. The 01BSUR antibody was obtained from Novartis under MTA that does not allow redistribution. Deidentified survey data is available on request from the authors. All other data are available in the main text or supplemental materials.

## SUPPLEMENTAL FIGURES

**Supplemental Table 1.**
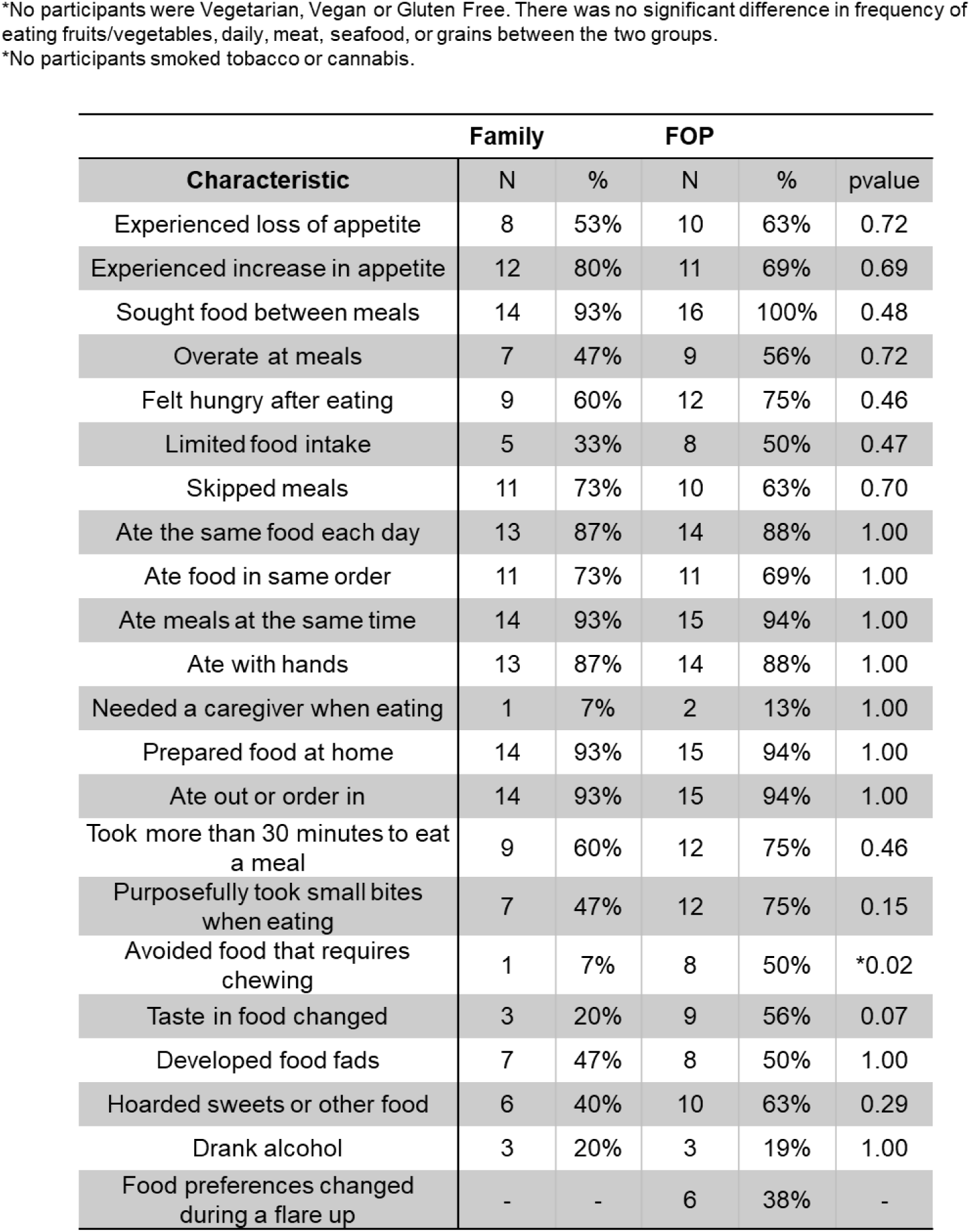
Comparison of self-reported food preferences and eating behaviors between individuals with FOP and family controls. Data are presented as counts (N) and percentages (%) for each group. Survey responses were dichotomized, and differences in frequency patterns of categorical variables were assessed using Fisher’s exact tests in Stata Statistics (College Station, TX; version 16). A statistically significant difference was observed only for avoidance of foods requiring chewing (*p* = 0.02), which was more common in individuals with FOP. No other eating behaviors differed significantly between groups. No participants reported vegetarian, vegan, or gluten-free diets, and no differences were observed in general dietary patterns (fruit/vegetable, meat, seafood, or grain consumption). No participants reported tobacco or cannabis use.

**Supplemental Table 2.** Comparison of self-reported gastrointestinal (GI) symptoms between individuals with FOP and family control participants. Data are presented as counts (N) and percentages (%) for each group, with total GI symptoms additionally reported as mean ± standard deviation (SD). Survey responses were dichotomized, and differences in frequency patterns of categorical variables were assessed using Fisher’s exact tests in Stata Statistics (College Station, TX; version 16). Vomiting (*p* = 0.01) and vomiting until the stomach was empty (*p* = 0.04) were significantly more frequent in individuals with FOP compared to family controls. All other GI symptoms were not significantly different between groups.

| Characteristic | Family |  | FOP |  | pvalue |
| --- | --- | --- | --- | --- | --- |
|  | N | % | N | % |  |
| Total GI Symptoms Reported (mean, sd) | 13 | 9.42 | 18.44 | 10.33 | 0.14 |
| Re-swallow food that came back into throat | 6 | 40% | 6 | 38% | 1.00 |
| Excess saliva in mouth | 10 | 67% | 9 | 56% | 0.72 |
| Burning in throat | 6 | 40% | 9 | 56% | 0.48 |
| Hiccups | 11 | 73% | 14 | 88% | 0.39 |
| Lump in throat | 6 | 40% | 7 | 44% | 1.00 |
| Food or liquid came back up into throat or mouth without vomiting | 4 | 27% | 6 | 38% | 0.70 |
| Burp | 15 | 100% | 15 | 94% | 1.00 |
| Substernal burning | 6 | 40% | 6 | 38% | 1.00 |
| Food stuck in throat while eating | 5 | 33% | 8 | 50% | 0.47 |
| Pain in chest when swallowing food | 5 | 33% | 8 | 50% | 0.47 |
| Difficulty swallowing solid foods | 3 | 20% | 6 | 38% | 0.43 |
| Difficulty swallowing soft foods | 1 | 7% | 2 | 13% | 1.00 |
| Difficulty swallowing liquids | 3 | 20% | 3 | 19% | 1.00 |
| Difficulty swallowing pills | 3 | 20% | 9 | 56% | 0.07 |
| Loose or watery stools | 10 | 67% | 12 | 75% | 0.70 |
| Felt need to use the bathroom right away or else would have an accident | 9 | 60% | 9 | 56% | 1.00 |
| Accident because could not make it to the bathroom in time | 1 | 7% | 6 | 38% | 0.08 |
| Leak stool or soil in underwear | 1 | 7% | 6 | 38% | 0.08 |
| Pass gas but stool or urine came out instead | 1 | 7% | 5 | 31% | 0.17 |
| Strain while going to the bathroom | 9 | 60% | 12 | 75% | 0.46 |
| Hard or lumpy stool | 8 | 53% | 13 | 81% | 0.14 |
| Pain while trying to pass a bowel movement | 7 | 47% | 9 | 56% | 0.72 |
| Need to use the bathroom after just went | 7 | 47% | 8 | 50% | 1.00 |
| Use finger or toilet paper to get out stool | 2 | 13% | 5 | 31% | 0.39 |
| Nausea | 7 | 47% | 12 | 75% | 0.15 |
| Know would have nausea before it happened | 4 | 27% | 5 | 31% | 1.00 |
| Vomit | 2 | 13% | 10 | 63% | *0.01 |
| Vomit until stomach empty | 1 | 7% | 7 | 44% | *0.04 |
| Swollen belly | 3 | 20% | 6 | 38% | 0.43 |
| Bloated | 6 | 40% | 8 | 50% | 0.72 |
| Bowel sounds when not hungry | 5 | 33% | 7 | 44% | 0.72 |
| Flatulence | 14 | 93% | 16 | 100% | 0.48 |
| Weight fluctuate by more than 5-10 lbs | 0 | 0% | 4 | 25% | 0.10 |
| Belly pain | 6 | 40% | 10 | 63% | 0.29 |
| Belly discomfort | 6 | 40% | 11 | 69% | 0.16 |
| Discomfort interferes with ADLs | 2 | 13% | 6 | 38% | 0.22 |

**Supplemental Table 3.** Differential abundance of bacterial taxa in individuals with FOP compared to healthy sibling controls using paired analysis. Differential abundance was assessed using DESeq2 (v1.24.0) with FDR-adjusted p-values after filtering low-abundance features. Symbols indicate directionality of change in FOP relative to controls (+, increased; −, decreased). At the phylum level, Firmicutes were increased and Bacteroidetes were decreased in FOP. At the species level, multiple taxa were differentially abundant, including decreased *Alistipes* spp., *Prevotella copri*, and *Veillonella* spp., and increased *Clostridium innocuum*, *Streptococcus mitis*, and *Turicibacter* spp.

Paired Analysis
| Phylum | FOP vs. WT | P value |
| --- | --- | --- |
| Firmicutes | + | 0.047 |
| Bacteroidetes | - | 0.016 |

| Species | FOP vs WT | P value |
| --- | --- | --- |
| *Alistipes senegalensis* | - | 0.0469 |
| *Alistipes finelgoldii* | - | 0.0156 |
| *Alistipes. Sp. HGB5* | - | 0.0313 |
| *Lachnospiracea*<br>*bacterium_3_1_57FAA_*<br>*CT1* | - | 0.0469 |
| *Prevotella copri* | - | 0.0156 |
| *Veillonella_u_s.* | - | 0.0156 |
| *Clostridium innocuum* | + | 0.0156 |
| *Bacterium LF3* | + | 0.0156 |
| *Erysipelotrichaceae*<br>*bacterium 21_3* | + | 0.0156 |
| *Streptococcus mitis* | + | 0.036 |
| *Turicibacter sanguinis* | + | 0.036 |
| *Turicibacter_sp_HGF1* | + | 0.036 |

**Supplemental Table 4.** Literature-based annotation of bacterial species differentially abundant in individuals with FOP compared to sibling controls. Directionality of change in FOP relative to controls is indicated (+, increased; −, decreased). Reported functional characteristics, classification as commensal or pathobiont, and potential beneficial or harmful roles are summarized based on published literature. References supporting each classification are listed. *Pathobiont in specific circumstances; † unknown in humans; ** harmful in certain medical conditions*.

| Phylum | Species | FOP vs. WT | Overview | Pathobiont or Commensal | Beneficial or Harmful | Reference |
| --- | --- | --- | --- | --- | --- | --- |
| **Bacillota**<br>(firmicutes) | *Veillonella u_s.* | - | Gram Negative | Commensal | Pathogenic classification (rare) | 1, 2 |
|  | *Streptococcus mitis* | + | Oral Commensal | Commensal, pathogenic in blood stream | Harmful | 3, 4, 5 |
|  | *Lachnospiraceae bacterium* | - | Protective | Pathobiont | Beneficial | 6, 7 |
|  | *Clostridium innocuum* | + | Emerging Pathogen | Commensal limited data | Harmful | 8, 9 |
|  | *Erysipelotrichaceae bacterium 21_3* | + | Pro inflammatory interhost variation | Commensal possible pathogen | Harmful Limited Data | 10, 11 |
|  | *Turcibacter sanguinis* | + | Gram Positive | Commensal | Beneficial | 12, 13 |
|  | *Turcibacter sp. HGF1* | + | Gut-Brain Axis | Commensal | Beneficial | 14 |
| **Bacteroidota**<br>(CFB group bacteria) | *Alistipes senegalensis* | - | Anti inflammatory | Commensal | Beneficial** | 15, 16 |
|  | *Alistipes finegoldii* | - | Anti inflammatory | Commensal | Both | 17 |
|  | *Alistipes sp. HGB5* | - | Unknown | Commensal | Harmful | 18, 19 |
|  | *Prevotella copri* | - | Inflammatory | Commensal | Diet specific & strain specific: Controversial results | 20 |

\*Pathobiont in specific circumstances
† Unknown in Humans
\*\*Harmful in certain medical conditions

**Supplemental Table 5.** Differentially abundant MetaCyc pathways in individuals with FOP compared to sibling controls identified by LEfSe analysis. Linear discriminant analysis (LDA) scores and Wilcoxon p-values are shown, with direction indicating enrichment in FOP or controls. LEfSe was performed using a Kruskal–Wallis α = 0.05, Wilcoxon α = 0.05, and an LDA score threshold of 2.0. Pathways with positive LDA scores are enriched in FOP, whereas negative scores indicate enrichment in controls.

| **MetaCyc pathway** | **LDA score** | **Wilcoxon p** | **Direction** |
| --- | --- | --- | --- |
| ppGpp biosynthesis | 2.59 | 0.03 | ↑ FOP |
| TCA cycle I (prokaryotic) | 2.85 | 0.03 | ↑ FOP |
| TCA cycle II (plants & fungi) | 2.79 | 0.05 | ↑ FOP |
| gluconeogenesis I | 2.86 | 0.02 | ↑ FOP |
| gondoate biosynthesis (anaerobic) | 3.20 | 0.02 | ↑ FOP |
| L-arginine biosynthesis (via N-acetyl-L-citrulline) | 2.68 | 0.02 | ↑ FOP |
| thiamin salvage II | 3.15 | 0.02 | ↑ FOP |
| purine ribonucleosides degradation | -3.21 | 0.03 | ↑ WT |
| D-galactose degradation V (Leloir pathway) | -3.01 | 0.02 | ↑ WT |
| galactose degradation I (Leloir pathway) | -3.01 | 0.02 | ↑ WT |
| stachyose degradation | -3.05 | 0.02 | ↑ WT |
| tetrapyrrole biosynthesis I (from glutamate) | -2.91 | 0.02 | ↑ WT |
| seleno-amino acid biosynthesis | -2.83 | 0.03 | ↑ WT |

**Supplemental Table 6.** Differentially abundant bacterial species in FOP mice compared to WT controls with literature-based functional annotation. Differential abundance was assessed using DESeq2 with FDR-adjusted p-values after filtering taxa present in >30% of samples. Directionality of change in FOP relative to controls is indicated (+, increased; −, decreased). Functional characteristics, classification as commensal or pathobiont, and potential beneficial or harmful roles are summarized from published literature. *Pathobiont in specific circumstances; † unknown in humans*.

| Phylum | Species | FOP vs. WT | Overview | Pathobiont or Commensal | Beneficial or Harmful | Reference |
| --- | --- | --- | --- | --- | --- | --- |
| **Actinomycetota**<br>(actinobacteria) | *Adlercreutzia equolifaciens* | + | Anti-inflammatory | Commensal | Beneficial | 21, 22 |
|  | *Aldercreutzia hattorii* | + | Gram-positive | Commensal | Beneficial | 23, 24 |
| **Bacillota**<br>(firmicutes) | *Flavonifractor plautii* | - | Anti Inflammatory | Commensal* | Beneficial | 25, 26 |
|  | *Acutalibacter muris* | - | Gram-Positive | Commensal† | No clinically relevant effects | 27, 28 |
|  | *Clostridium Scindens* | - | Gram-Positive | Commensal* | Beneficial | 29 |
|  | *Simiaoa sunii* | - | Gram Negative | Commensal | Unknown | 30 |
|  | *Blautia obeum* | + | Reduced inflammation | Commensal | Beneficial | 31, 32 |
| **Bacteroidota**<br>(CFB group bacteria) | *Alistipes finegoldii* | + | Immunotherapy response implications | Pathobiont* | Harmful | 33, 34 |
|  | *Bacteroides thetaiotaomicron* | + | Anti inflammatory properties | Commensal | Beneficial | 35, 36 |
|  | *Parabacteroides johnsonii* | + | Anti inflammatory | Commensal | Beneficial | 37, 38 |
|  | *Muribaculum gordoncarteri* | + | Anti inflammatory | Commensal | Beneficial | 39, 40 |
| **Campylobacterota**<br>(e-proteobacteria) | *Helicobacter cinaedi* | - | Pro inflammatory Gram-Negative | Pathobiont | Harmful | 21, 22 |
|  | *Helicobacter hepaticus* | - | Pro inflammatory | Pathobiont | Harmful | 23 |
|  | *Helicobacter typhlonius* | - | Pro inflammatory | Pathobiont | Harmful | 24 |

\*Pathobiont in specific circumstances
† Unknown in Humans

**Supplemental Table 7.** Summary of cytokine changes across experimental comparisons in FOP mice and related datasets. Arrows indicate direction of change (↑ higher, ↓ lower), NS indicates no significant difference, and “–” indicates not tested. Comparisons include genotype (FOP vs WT), effects of GMA, and validation from previously published serum and iPSC-derived macrophage (iMAC) studies.

| Cytokine | FOP vs WT | FOP GMA vs WT | FOP GMA vs FOP | WT GMA vs WT | FOP GMA vs WT GMA | Serum FOP vs WT<sup>45</sup> | IMAC FOP vs WT<sup>46</sup> |
| --- | --- | --- | --- | --- | --- | --- | --- |
| IL-1α | ↑ | NS | ↓ | NS | NS | - | ↑ |
| IL-2 | ↑ | NS | NS | ↓ | NS | - | - |
| MIP-1α | ↑ | ↑ | NS | NS | NS | ↑ | ↑ |
| IL-11 | ↑ | ↑ | NS | NS | NS | - | - |
| TIMP-1 | ↑ | NS | NS | ↑ | NS | - | - |
| 6Ckine/<br>Exodus 2 | ↓ | NS | NS | ↑ | NS | - | - |
| fractalkine | ↓ | NS | NS | ↓ | NS | ↑ | - |
| G-CSF | NS | ↑ | NS | ↓ | NS | ↑ | - |
| eotaxin | NS | NS | ↓ | NS | NS | ↑ | - |
| IL-6 | NS | NS | ↓ | NS | NS | - | ↑ |
| MIP-1β | NS | NS | ↓ | NS | ↑ | ↑ | ↑ |

↑ = higher
↓ = lower
NS = no significant difference
- = not tested

**Supplemental Table 8.** Clinical characteristics of study participants, including individuals with fibrodysplasia ossificans progressiva (FOP) and related family controls. Data include sex, age, FOP status, cumulative joint involvement scale (CAJIS), microbiome pairing, medical history, and current medications. Clinical and survey data were collected using REDCap-based questionnaires adapted from validated instruments, and CAJIS scores were obtained via in-person or video assessment. One subject was unable to complete the CAJIS assessment.

| ID | Sex | Age | FOP Status | CAJIS | Microbiome | Medical History | Medications |
| --- | --- | --- | --- | --- | --- | --- | --- |
| 16 | F | 12 | FOP | 18 | Sibling Pair 5 | Anxiety, seizure/syncope, hearing loss, restricted lung volume, low weight | acetaminophen, celecoxib, fluoxetine, lorazepam, ibuprofen |
| 17 | M | 15 | Sibling |  | Sibling Pair 5 | None | None |
| 14 | M | 14 | FOP | 17 |  | Childhood seizure, peripheral neuropathy, hearing loss, asthma, restrictive lung disease | calcium carbonate, ceterizine, vitamin D3, ibuprofen, ketoprofen topical, meloxicam, lansoprazole, |
| 15 | F | 10 | Family |  |  | None | None |
| 20 | F | 14 | FOP | 1 | Sibling Pair 1 | Asthma | Albuterol Inhaler |
| 21 | M | 15 | Sibling |  | Sibling Pair 1 | None | None |
| 1 | F | 9 | FOP | 5 | Sibling Pair 2 | Restricted chest expansion, hearing loss | Celecoxib, Cromolyn, Montelukast, Apigenin |
| 2 | M | 6 | Sibling |  | Sibling Pair 2 | None | None |
| 22 | M | 11 | FOP | 8 | Sibling Pair 3 | Hearing Loss, restricted chest expansion | Cromolyn, Montelukast, Ranitidine |
| 23 | F | 13 | Sibling |  | Sibling Pair 3 | None | None |
| 8 | M | 11 | FOP | Missing |  |  | celecoxib, montelukast, topical triamcinolone, palovarotene |
| 9 | M | 9 | Sibling |  |  | None | None |
| 12 | M | 6 | FOP | 12 | Sibling Pair 4 | Hearing loss, asthma, muscle spasms, headaches, cognitive impairment, restricted chest expansion | Celecoxib, Beclomethasone inhaler, Vitamin D, ipatropium inhaler, montelukast, methylphenidate |
| 13 | F | 14 | Sibling |  | Sibling Pair 4 | None | None |
| 24 | M | 11 | FOP | 6 |  | Pematurity | None |
| 25 | F | 9 | Sibling |  |  | None | None |
| 6 | F | 15 | FOP | 5 |  | None | cromolyn, iron sulfate, montelukast |
| 7 | F | 21 | Full sibling |  |  | Asthma | iron supplement, albuterol |
| 3 | M | 11 | FOP | 5 |  | Hearing loss | Montelukast, Miralax; multivitamin, calcium |
| 4 | F | 6 | Sibling |  |  | None | Multivitamin, calcium |
| 5 | F | 15 | Sibling |  |  | None | Multivitamin, calcium |
| 26 | M | 9 | FOP | 2 | Sibling Pair 6 | Ear infections, heart failure | Celecoxib, cromolyn, montelukast, apigenin, fish oil |
| 27 | M | 11 | Sibling |  | Sibling Pair 6 | Prematurity | fluoxetine, adderall, guaifacine, folic acid, melatonin |
| 40 | F | 30 | FOP | 20 |  | Hearing loss, restricted chest expansion, hyperlipidemia | Seasonque, cromolyn, palovarotene, acidophilus probiotic, biotin, vitamin D, Calcium |
| 38 | F | 31 | FOP | 8 |  | Hearing loss, ear infections, prior fracture | Gastrocom, singulare, celebrex sprintek, |
| 37 | M | 33 | Non-cohabitating sibling |  |  | None | None |
| 32 | M | 11 | FOP | 10 | Sibling Pair 7 | Hearing loss, restricted chest expansion, irritable bowel disease, pediatric pyloric stenosis | Omega 3, vitamin D |
| 33 | M | 13 | Sibling |  | Sibling Pair 7 | Cranial Gorham-Stout | Levothyroxine, omega 3, vitamin D |
| 34 | M | 49 | FOP | 22 |  | Hearing loss, asthma, locked jaw, muscle spasms, prior hernias | Claritin, duloxetine, acetaminophen, prilosec, cymbalta, brioalpipt inhaper |
| 35 | F | 52 | Wife |  |  | Cerebral Palsey, hemorrhoids, osteoarthritis | Buspar, prilosec, claritin, fiber, B12 |
| 41 | M | 19 | FOP | 4 |  | Hearing loss, ear infections, arrhythmias | Palovarotene |

**Supplemental Figure 1.**
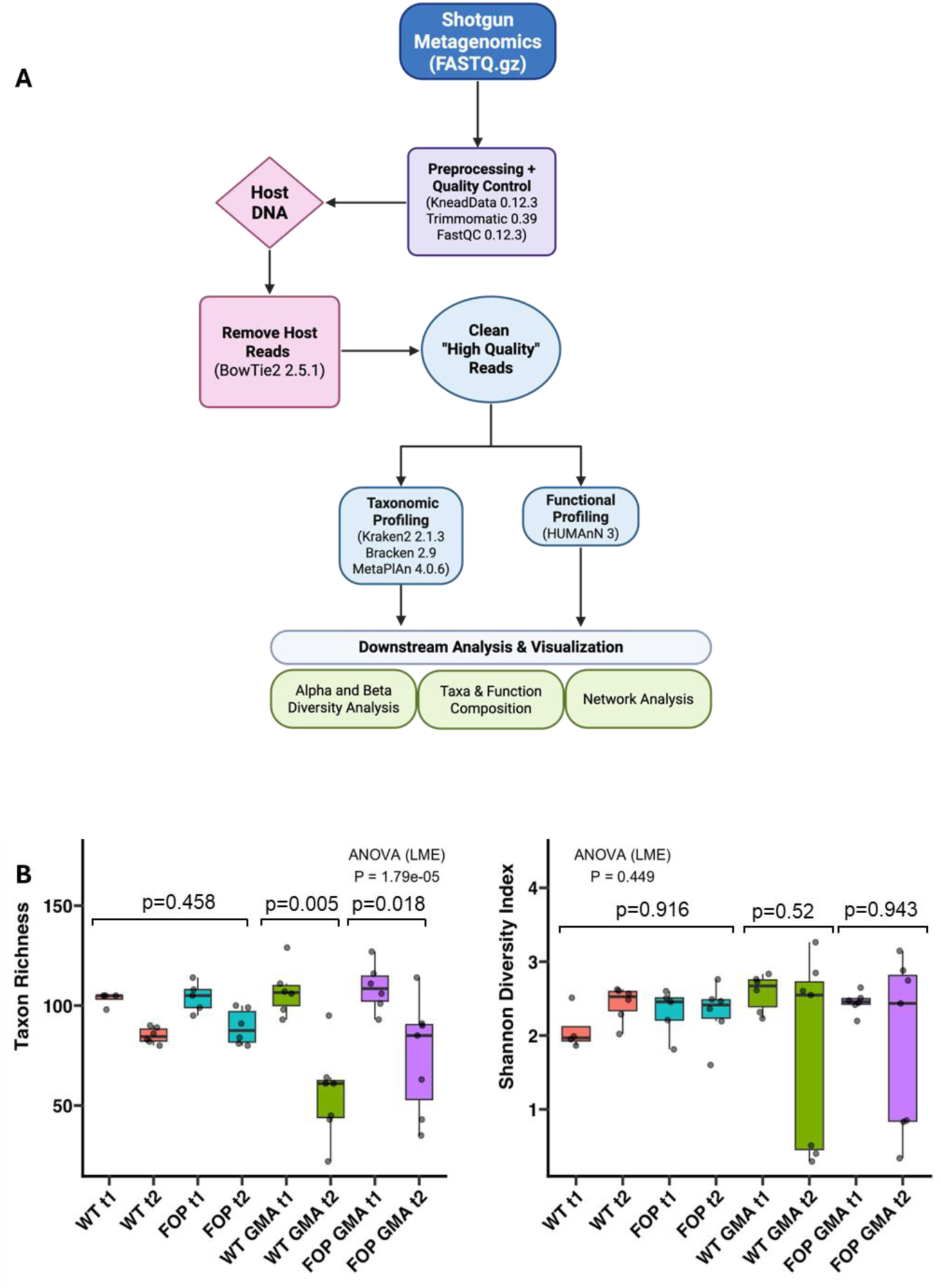
Mouse metagenomics workflow and alpha diversity analysis. **(A)** Overview of the shotgun metagenomics pipeline. Fecal DNA was sequenced (Illumina NovaSeq 6000, 2×150 bp), followed by quality control and trimming (FastQC, KneadData/Trimmomatic), host read removal (Bowtie2), and generation of high-quality microbial reads. Taxonomic profiling was performed using Kraken2/Bracken (validated with MetaPhlAn), and functional profiling was conducted using HUMAnN3. **(B)** Alpha diversity across experimental groups, baseline (t1) and post-treatment (t2). Taxon richness and Shannon diversity were calculated at the species level from normalized count data. Group differences were assessed using linear mixed-effects models (ANOVA) with MouseID as a random effect, and p-values are indicated. A significant difference in taxon richness was observed between WT GMA t1 and WT GMA t2 groups (p = 0.005) as well as FOP GMA t1 and FOP GMA t2 (p = 0.018), while no significant differences were observed in Shannon diversity.

**Supplemental Figure 2.**
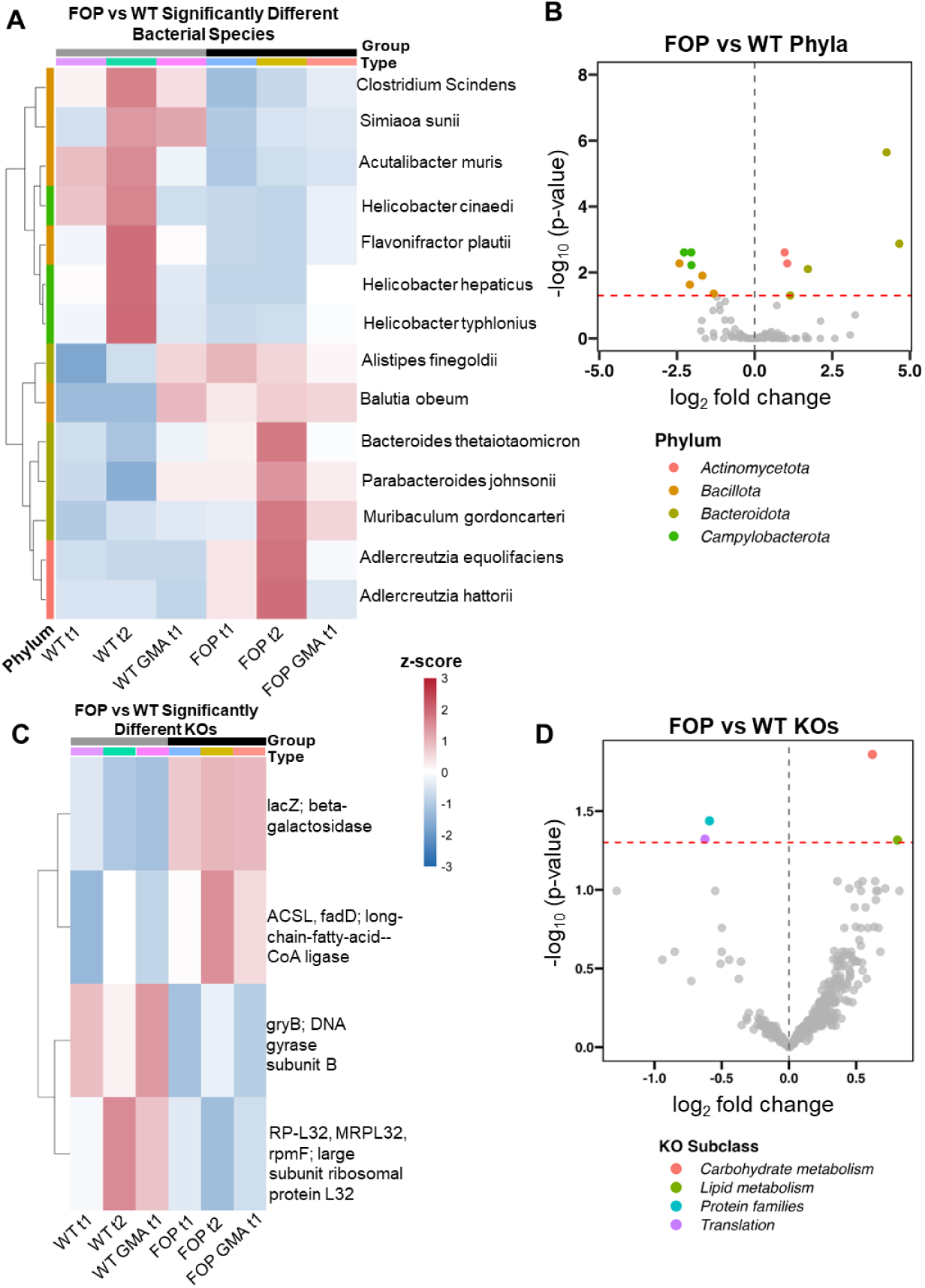
Differential taxonomic and functional profiles in FOP mice. **(A)** Heatmap of significantly different bacterial species between FOP and WT mice, +/-GMA. Species-level counts were generated from Kraken2/Bracken outputs, normalized using DESeq2 size factors, log₁₀-transformed, and standardized by row-wise z-scoring. **(B)** Volcano plot of phylum-level differential abundance (log₂ fold change vs. −log₁₀ adjusted p-value). Differential abundance was assessed using DESeq2 with Wald tests and Benjamini–Hochberg FDR correction (padj < 0.05). Positive log₂ fold change values indicate taxa increased in FOP, whereas negative values indicate taxa decreased in FOP relative to WT. **(C)** Heatmap of significantly different KEGG orthologs (KOs) between FOP and WT mice, +/-GMA. KO abundances were derived from HUMAnN3, normalized to copies per million, log₁₀-transformed, and z-scored by row. **(D)** Volcano plot of KO-level differential abundance (log₂ fold change vs. −log₁₀ FDR-adjusted p-value). Positive log₂ fold change values indicate features increased in FOP, whereas negative values indicate features decreased in FOP relative to WT; significant features (padj < 0.05) are highlighted and colored by functional subclass.

**Supplemental Figure 3.**
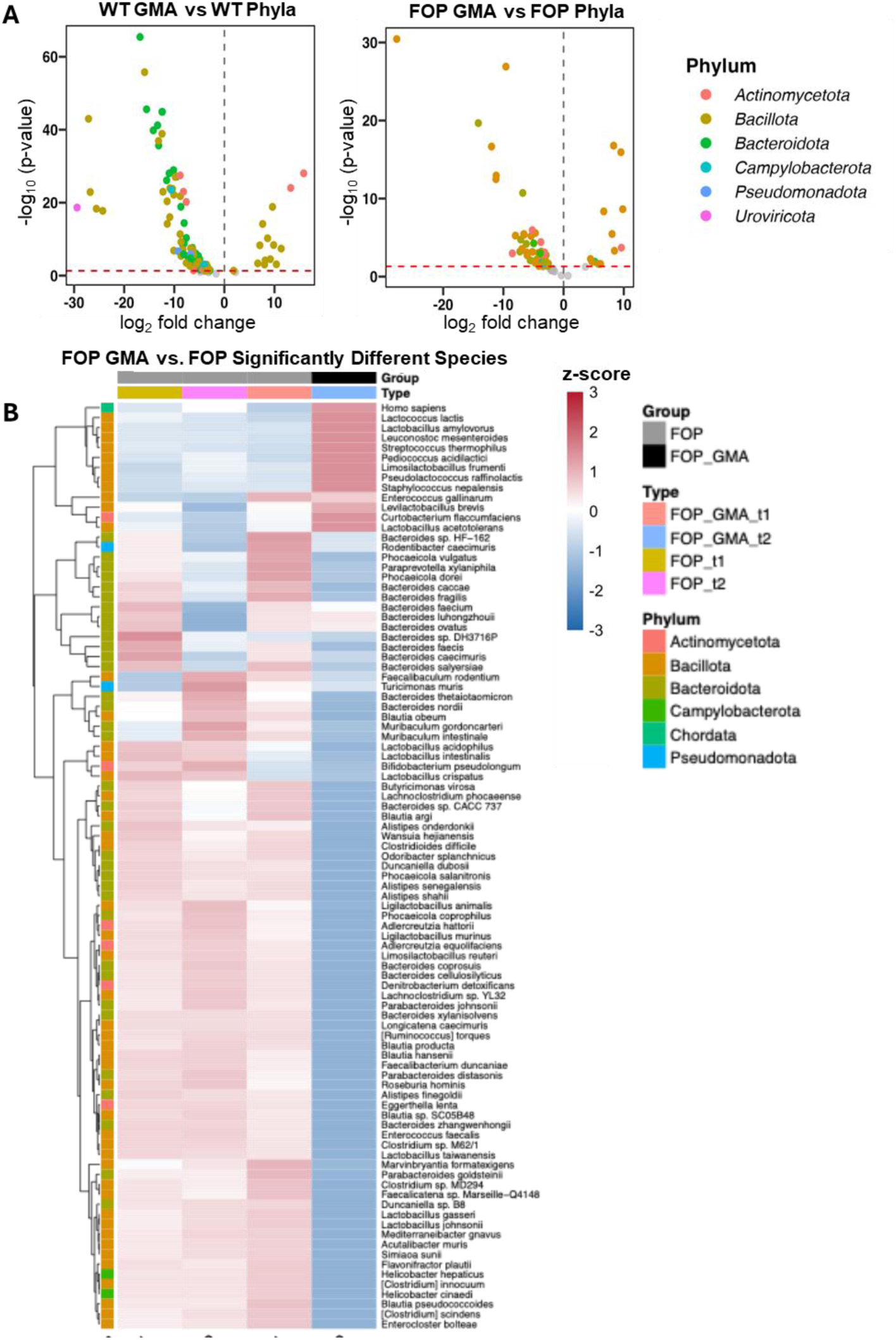
Significantly different bacterial taxa following GMA. **(A)** Volcano plots showing phylum-level differential abundance for WT GMA vs WT and FOP GMA vs FOP (log₂ fold change vs. −log₁₀ adjusted p-value). Differential abundance was assessed using DESeq2 with Wald tests and Benjamini–Hochberg FDR correction (padj < 0.05). Positive log₂ fold change values indicate taxa increased following GMA, whereas negative values indicate taxa decreased following GMA. **(B)** Heatmap of significantly different bacterial species between FOP GMA and FOP mice, baseline (t1) and post-treatment (t2). Species-level counts were generated from Kraken2/Bracken outputs, normalized using DESeq2 size factors, log₁₀-transformed, and standardized by row-wise z-scoring. Samples are grouped by genotype and timepoint.

**Supplemental Figure 4.**
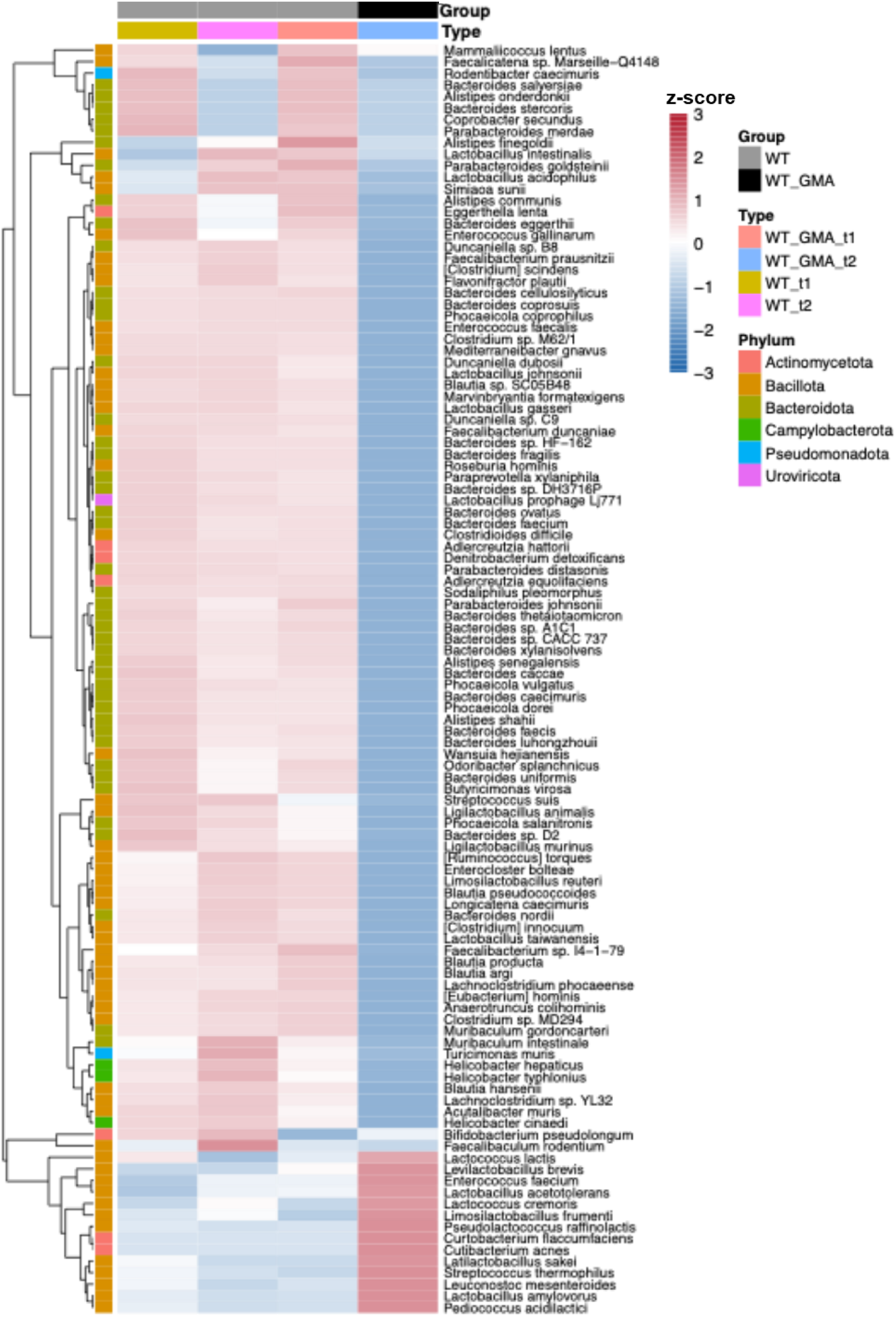
Significantly different species following GMA in WT mice. Heatmap of significantly different bacterial species between WT GMA and WT mice, baseline (t1) and post-treatment (t2). Species-level counts were generated from Kraken2/Bracken outputs, normalized using DESeq2 size factors, log₁₀-transformed, and standardized by row-wise z-scoring. Differential abundance was assessed using DESeq2 with Wald tests and Benjamini–Hochberg FDR correction (padj < 0.05). Samples are grouped by treatment and timepoint.

**Supplemental Figure 5.**
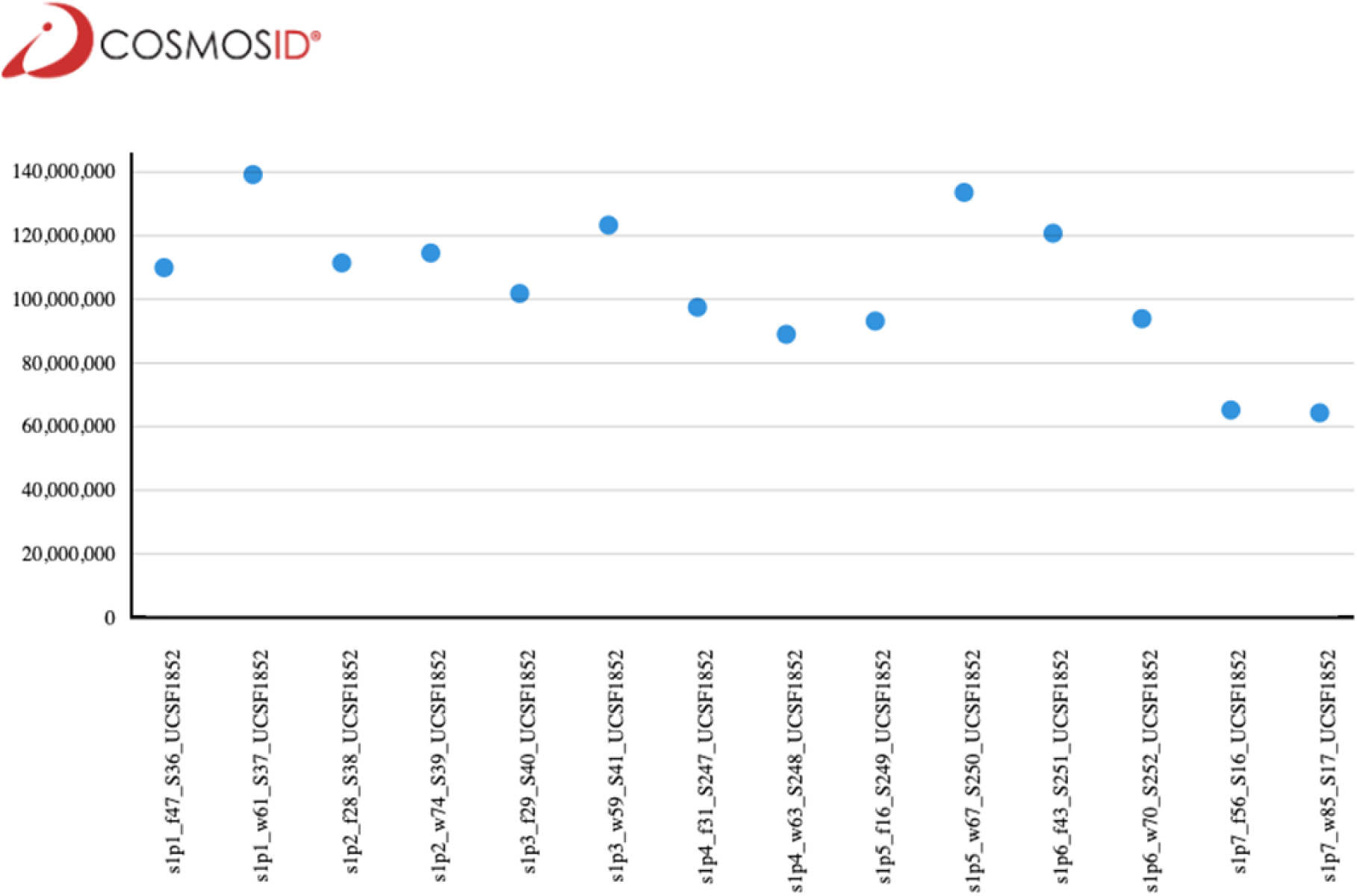
Shotgun metagenomic sequencing depth across human stool samples. Human fecal samples were collected using home collection kits, stored at −20 °C prior to shipment, and archived at −80 °C upon receipt. DNA was extracted using a bead-beating protocol and sequenced on Illumina HiSeq 4000 or NovaSeq 6000 platforms (150-bp paired-end reads). Sequencing depth ranged from ∼65–140 million reads per sample, with no major outliers, indicating consistent and sufficient coverage for downstream taxonomic and functional analyses.

**Supplemental Figure 6.**
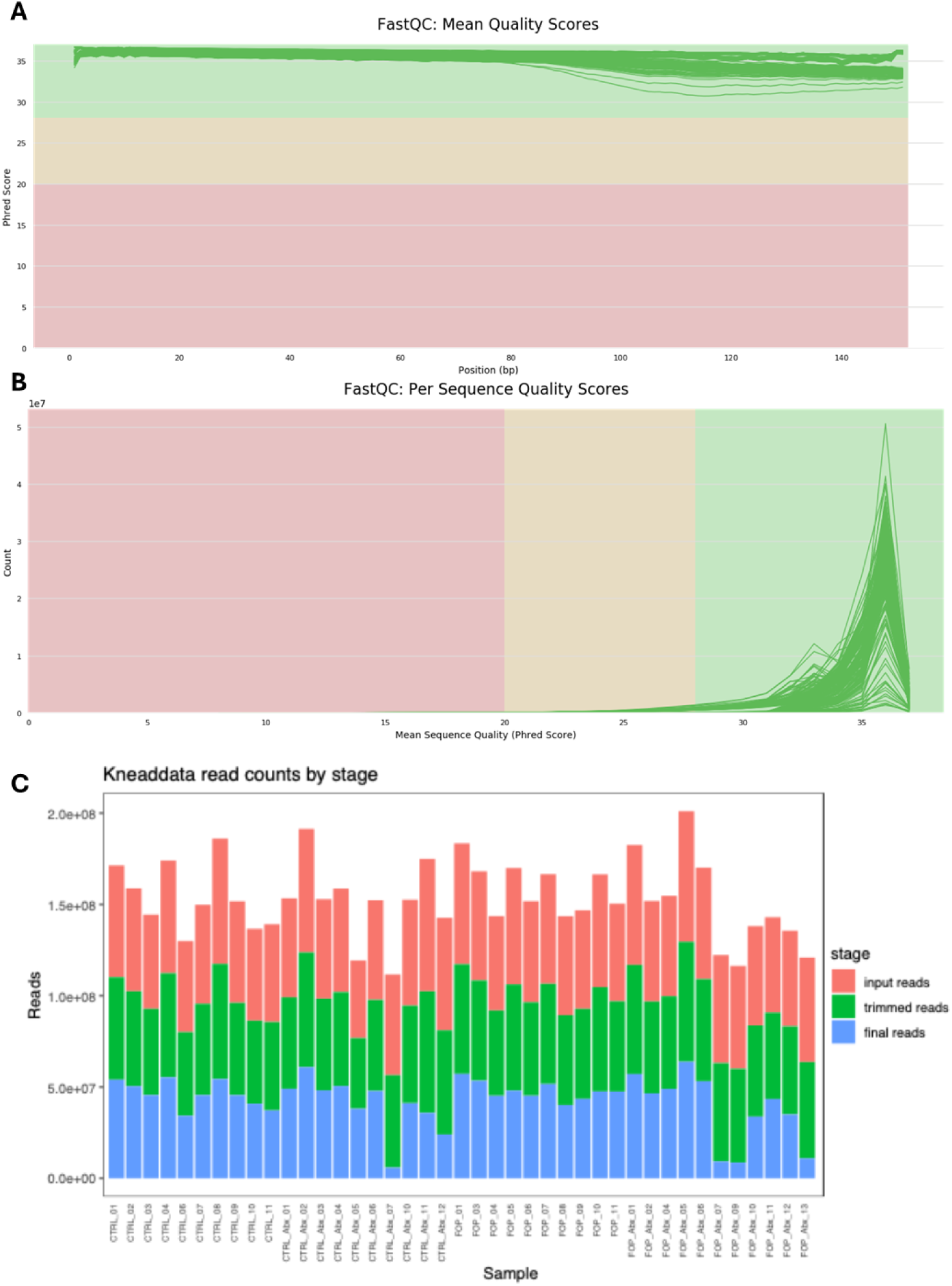
Mouse shotgun metagenomic sequencing quality control. **(A)** Mean per-base sequence quality scores generated using FastQC (v0.12.1) across all samples, demonstrating consistently high base-call quality (Phred scores >30). **(B)** Per-sequence quality score distributions from FastQC. **(C)** Read counts across preprocessing stages generated using KneadData (v0.12.3), which incorporates Trimmomatic (v0.39) for adapter trimming and quality filtering and Bowtie2 (v2.5.1) for host read removal. Reads were filtered using a sliding 4-bp window (minimum Phred score of 20), and reads <50 bp or containing adapter contamination were removed. The reduction in reads from input to trimmed to final reflects quality filtering and host depletion, with substantial retention of high-quality microbial reads for downstream analysis.

**Supplemental Figure 7.**
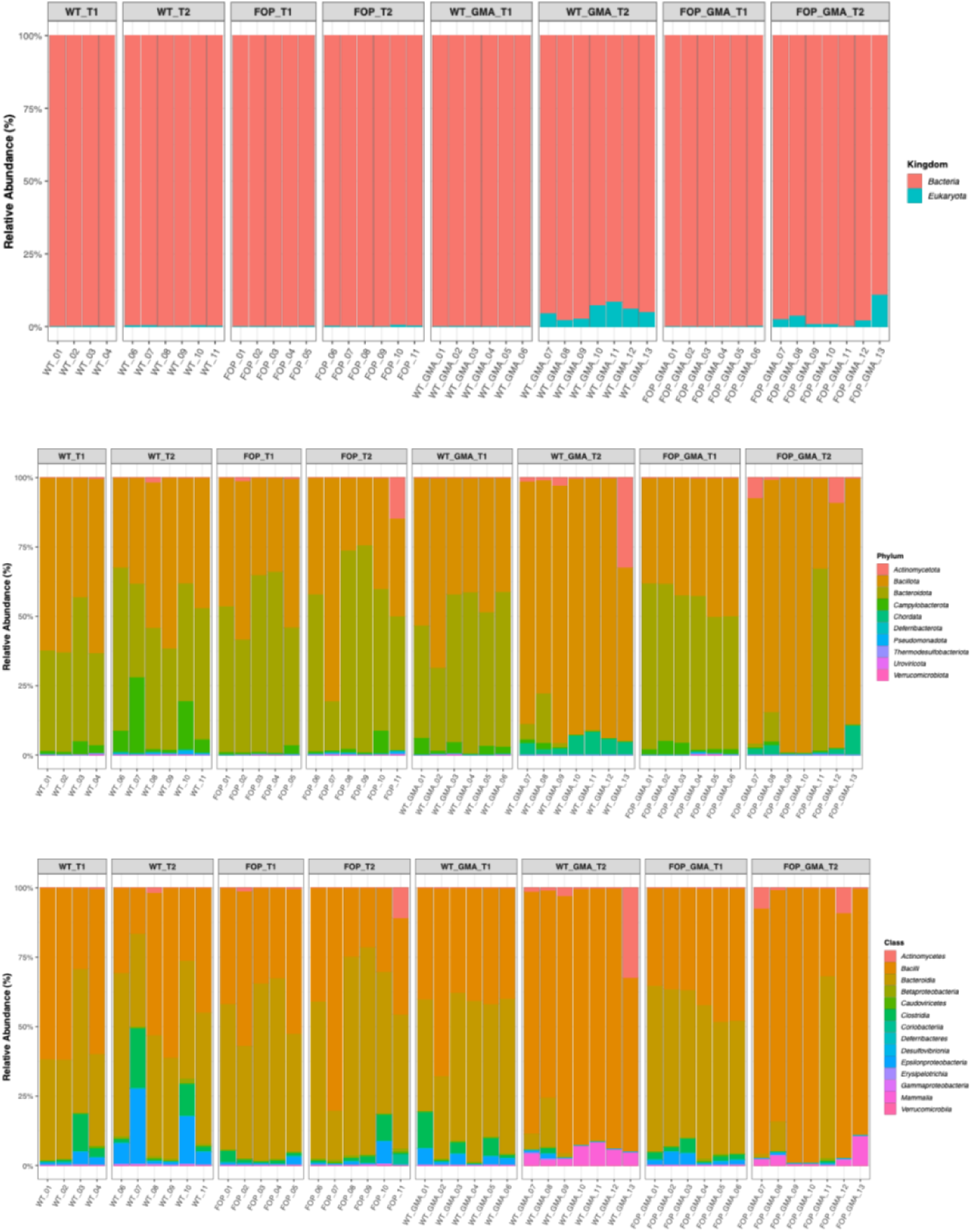
Relative abundance of microbial taxa across experimental groups. Stacked bar plots showing relative abundance of microbial communities across WT, FOP, and GMA groups at baseline (t1) and post-treatment (t2). Taxonomic profiles were generated using Kraken2 (v2.1.3) with species-level abundance estimates refined by Bracken (v2.9), with validation using MetaPhlAn 4.0.6. Taxa present in <10% of samples or at very low abundance were filtered prior to analysis. Relative abundances were calculated on a per-sample basis and aggregated at the kingdom (top), phylum (middle), and class (bottom) levels for visualization.

